# Increased immunogen valency improves the maturation of vaccine-elicited HIV-1 VRC01-class antibodies

**DOI:** 10.1101/2025.03.13.642975

**Authors:** Parul Agrawal, Arineh Khechaduri, Kelsey R. Salladay, Anna MacCamy, Duncan K. Ralph, Andrew Riker, Andrew B. Stuart, Latha Kallur Siddaramaiah, Xiaoying Shen, Frederick A. Matsen, David Montefiori, Leonidas Stamatatos

**Affiliations:** Vaccine and Infectious Disease Division, Fred Hutchinson Cancer Center, Seattle, WA, USA; Computational Biology Program, Fred Hutchinson Cancer Center, Seattle, WA, USA; Translational Science and Therapeutics Division, Fred Hutchinson Cancer Center, Seattle, WA, USA; Division of Surgical Sciences, Duke University Medical Center, Durham, NC, USA; Department of Global Health, University of Washington, Seattle, WA, USA; Howard Hughes Medical Institute, Computational Biology Program, Fred Hutchinson Cancer Center, Seattle, WA, USA; Department of Genome Sciences, University of Washington, Seattle, WA, USA; Department of Statistics, University of Washington, Seattle, WA, USA

## Abstract

Antibodies belonging to the VRC01-class display broad and potent neutralizing activities and have been isolated from several people living with HIV (PLWH). A member of that class, monoclonal antibody VRC01, was shown to reduce HIV-acquisition in two phase 2b efficacy trials. VRC01-class antibodies are therefore expected to be a key component of an effective HIV-1 vaccine. In contrast to the VRC01-class antibodies that are highly mutated, their unmutated forms do not engage HIV-1 envelope (Env) and do not display neutralizing activities. Hence, specifically modified Env-derived proteins have been designed to engage the unmutated forms of VRC01-class antibodies, and to activate the corresponding naïve B cells. Selected heterologous Env must then be used as boost immunogens to guide the proper maturation of these elicited VRC01-class antibodies. Here we examined whether and how the valency of the prime and boost immunogens influences VRC01-class antibody-maturation. Our findings indicate that, indeed the valency of the immunogen affects the maturation of elicited antibody responses by preferentially selecting VRC01-class antibodies that have accumulated somatic mutations present in broadly neutralizing VRC01-class antibodies isolated from PLWH. As a result, antibodies isolated from animals immunized with the higher valency immunogens display broader Env cross-binding properties and improved neutralizing potentials than those isolated from animals immunized with the lower valency immunogens. Our results are relevant to current and upcoming phase 1 clinical trials that evaluate the ability of novel immunogens aiming to elicit cross-reactive VRC01-class antibody responses.

**AUTHOR SUMMARY:** PA performed ELISA, B cell sorting, BCR sequencing analysis, oversaw the analysis of all immunochemical assays, interpreted the results and wrote the manuscript; AK processed tissues from immunized animals, performed B cell staining, and generated mAbs; KRS performed BLI and BCR sequencing; AM performed BCR sequencing; DR and FAM performed phylogenetic analysis and contributed to writing the manuscript; AR performed ELISA and processed tissues from immunized animals, ABS expressed and purified recombinant Envs; LKS expressed and purified mVRC01 mAb; XS and DM performed neutralization assays; LS conceived and oversaw the study and wrote the manuscript.

## INTRODUCTION

An estimated 39.9 million people were living with the human immunodeficiency virus (HIV) at the end of 2023 and an estimated 1.3 million new infections occurred in 2023 globally (WHO), despite the development of effective antiretroviral drugs (Nachega et al. 2023; hivinfo.nih.gov 2024). An effective HIV vaccine is therefore needed for the significant reduction of the number of new infections occurring. Such a vaccine would elicit diverse immune responses, including broadly neutralizing antibodies (bnAbs).

bnAbs have been isolated from people living with HIV (PLWH), and their structures as well as those of their epitopes on the HIV-1 envelope (Env), have been well characterized (Burton and Hangartner 2016; Mascola and Haynes 2013; West et al. 2014; Haynes et al. 2023). bnAbs that recognize the same region of Env and share common genetic and structural features are grouped into ‘classes’ (Kwong and Mascola 2012), and one such class are the VRC01-class which recognize a conserved epitope within the CD4-binding site (CD4-BS) of the viral Env.

Their VHs are derived from the VH1-2*02 allele while their light chains (LCs) express 5 amino acid (5-aa) long CDRL3 domains (Scheid et al. 2011; Wu et al. 2011; Wu et al. 2015; West et al. 2012; Zhou et al. 2013; Zhou et al. 2015; Scharf et al. 2013; Scharf et al. 2016; Umotoy et al. 2019). They are among the most mutated bnAbs known (Klein et al. 2013) and can display up to 50 percent aa sequence divergence; yet they recognize their epitope on diverse Envs with similar angles of approach (Scheid et al. 2011; Wu et al. 2011; Zhou et al. 2015). VRC01-class bnAbs protect animals from experimental S/HIV infection (Balazs et al. 2014; Shingai et al. 2014) and one mAb of this class, VRC01, was shown to prevent HIV-1 acquisition from susceptible, circulating primary HIV-1 viruses, in two phase 2b efficacy trials (Corey et al. 2021). Thus, we expect VRC01-class bnAbs to be a component of the immune responses elicited by an effective HIV-1 vaccine.

Although VRC01-class bnAbs bind diverse Envs and potently neutralize HIV-1 viruses from different clades, their unmutated forms (germline ‘gl’) do not (Hoot et al. 2013; Jardine et al. 2013; McGuire et al. 2013; McGuire et al. 2014). As a result, B cells expressing glVRC01-class B cell receptors (BCRs) are not activated by diverse Env-derived immunogens (Hoot et al. 2013; Jardine et al. 2013; McGuire et al. 2013; McGuire et al. 2014). To overcome this lack of naïve B cell-activation, specifically modified Env-derived constructs have been designed (McGuire et al. 2016; Jardine et al. 2013; McGuire et al. 2013; Jardine et al. 2016; Medina-Ramirez et al. 2017); such constructs, commonly referred to as ‘germline-targeting’ immunogens (Stamatatos, Pancera, and McGuire 2017), bind the germline VRC01-class antibodies/BCR forms and activate the corresponding B cells (McGuire et al. 2016; Jardine et al. 2016; Dosenovic et al. 2015; Parks et al. 2019; Lin et al. 2020). A common, key feature of germline-targeting Env-derived immunogens is the elimination of the conserved N-linked glycosylation site (NLGS) at position 276 (N276), within the Loop D region of the gp120 subunit. The carbohydrates at N276 represent the major steric block for glVRC01-class antibodies (Zhou et al. 2013; McGuire et al. 2013; Borst et al. 2018), but during affinity maturation the antibodies accumulate specific mutations in their heavy chain (HC) and light chain (LC) that allow them to overcome this problem (Scheid et al. 2011; Wu et al. 2011; Wu et al. 2015; Zhou et al. 2013; Barnes et al. 2022; Bonsignori et al. 2018).

One such germline-targeting immunogen is 426c.Mod.Core (previously referred to as TM4ΔV1– 3; (McGuire et al. 2016)), derived from the clade C 426c virus. This immunogen efficiently activates naïve B cells expressing glVRC01-class BCRs in transgenic animal models expressing human glVRC01-class BCRs (Parks et al. 2019; Lin et al. 2020; Knudsen et al. 2022; Agrawal et al. 2024) and is currently evaluated in a phase 1 clinical trial HVTN301(ClinicalTrials.gov NCT05471076). While immunization of transgenic mice expressing human glVRC01-class-derived VH/VL genes with 426c.Mod.Core results in the activation and partial maturation (through the accumulation of somatic mutations) of naive B cells expressing glVRC01-class BCRs, heterologous Env boost immunizations are expected to be necessary for the further maturation of those BCRs towards their broadly neutralizing forms (Dosenovic et al. 2015; Parks et al. 2019; Knudsen et al. 2022; Tian et al. 2016; Briney et al. 2016; Chen et al. 2021; Agrawal et al. 2024). The Env-derived proteins used as ‘boosts’ are expected to express glycans at N276. In our previous studies, we have employed the heterologous HxB2.WT.Core (derived from clade B) as our first booster immunogen. HxB2.WT.Core Env by itself does not activate germline VRC01 B cells (Knudsen et al. 2022). Importantly, the VRC01 antibodies isolated following the HxB2.WT.Core Env can accommodate the N276-associated glycans and as a result they display higher Env binding affinities, and improved neutralizing potentials, as compared to the VRC01-like antibodies elicited by the 426c.Mod.Core immunization alone (Parks et al. 2019; Knudsen et al. 2022).

In previous experiments we employed two oligomerization forms of our germline-targeting immunogen 426c.Mod.Core, and the first boost immunogen HxB2.WT.Core: (a) as Ferritin-based nanoparticles (NP) (24meric) of these immunogens (Knudsen et al. 2022; Agrawal et al. 2024) and (b) as self-assembling NP form (5-7meric) based on the oligomerization motif of the C4b-binding protein (Parks et al. 2019; McGuire et al. 2016; Ogun et al. 2008; Hofmeyer et al. 2013). Antigen valency has multifaceted effects on B cell responses (Kato et al. 2020), but we have not yet examined whether and how the valencies of our prime and boost immunogens affect the activation and maturation of VRC01-class BCRs. Here, we compared the VRC01 B cell and antibody responses elicited by Ferritin-based and C4b-based NPs of 426c.Mod.Core, and of HxB2.WT.Core Envs. We employed the TLR7/8 agonist 3M-052-AF + Alum adjuvant, which induces potent antigen-specific immune responses in non-human primates, characterized by Th1 cellular responses; as well as long-lived antibody and plasma cell responses (Kasturi et al. 2020). In a recent phase 1 clinical evaluation, it was also shown to be more effective than other adjuvants in eliciting autologous tier 2 HIV-1 neutralizing responses (Hahn et al. 2024)(ClinicalTrials.gov NCT04177355).

## RESULTS

### The valency of 426c.Mod.Core affects the potency of elicited anti-CD4-BS antibodies

As wild type animal species, including mice and non-human primates, do not express orthologs of the human VH1-2*02 allele (West et al. 2012; Jardine et al. 2013; Vigdorovich et al. 2016), immunization studies aiming at the elicitation of VRC01-class antibodies, are performed in transgenic mouse models expressing human VRC01-related VH/VL genes (Dosenovic et al. 2015; Tian et al. 2016; Jardine et al. 2015; Caniels et al. 2023; Luo et al. 2023). Here, we utilized the glVRC01^HC^ mouse model that is heterozygous for the human inferred glHC of the VRC01 mAb (Jardine et al. 2015). The estimated frequency of naive B cells expressing potential glVRC01 BCRs in this mouse model is approximately 0.08% (compared to approximately ∼ 0.0002% in humans).

A single immunization with either NP form of 426c.Mod.Core, elicited robust autologous plasma antibody responses in all animals (Fig 1A), but the antibody titers were significantly higher (p=0.029; Mann-Whitney test) in animals immunized with the 426c.Mod.Core.Fer NP than with the 426c.Mod.Core.C4b NP (Fig 1C, red circles). Antibodies cross-reacting with the heterologous HxB2.WT.Core Env were generated by all animals immunized with either NP form (Fig 1B), with no significant differences in the titers of these antibodies between the two NP groups.

**Fig 1.**
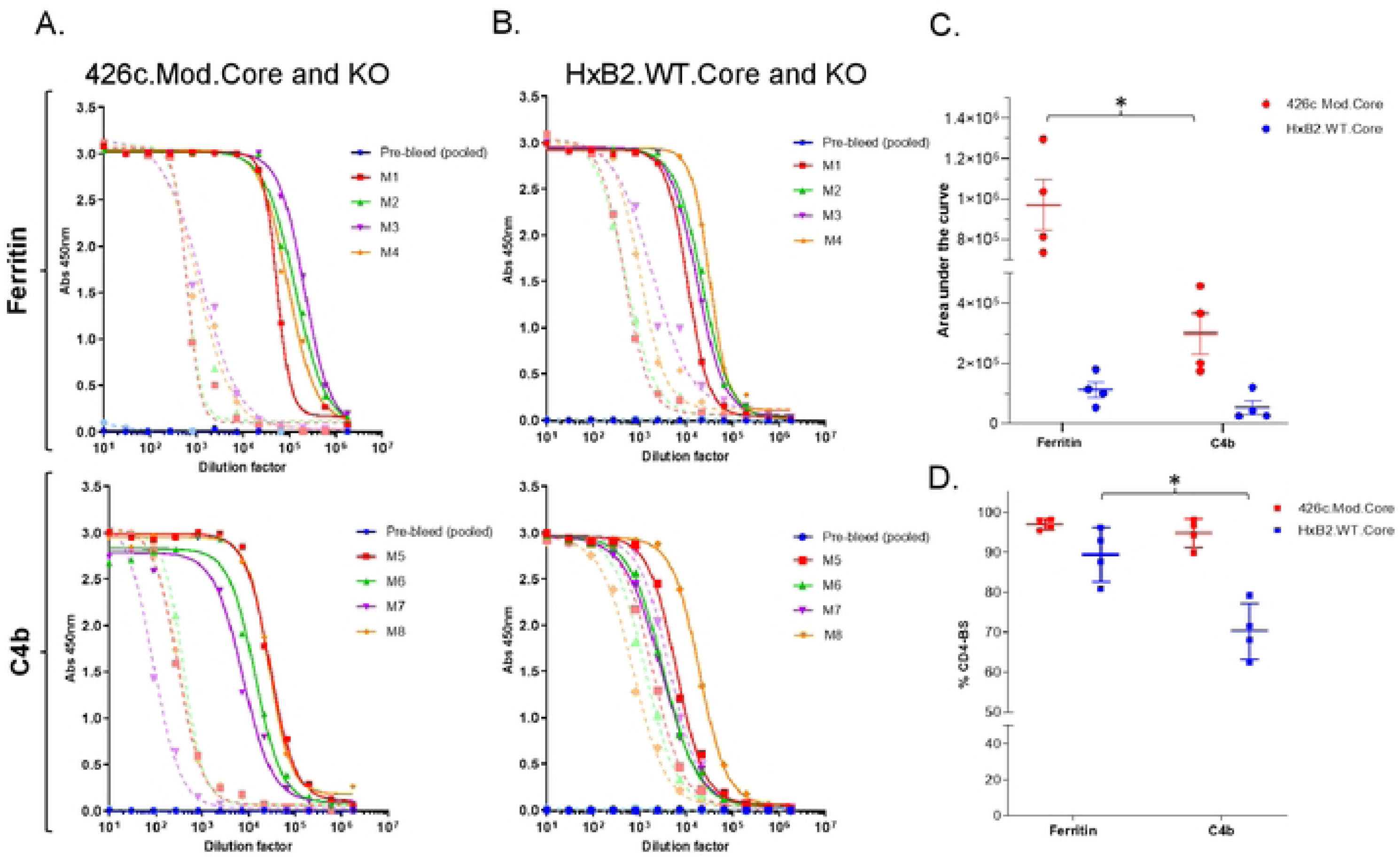
Antibody responses 2 weeks following the prime immunization. Mice (n = 4) were primed with adjuvanted 426c.Mod.Core-Fer or C4b NPs at week 0 and plasma at week 2 was assayed by ELISA. (A and B) Binding against 426.Mod.Core (solid lines), HxB2.WT.Core (solid lines), as well as their corresponding CD4-BS knock-out antigens (KO; dotted lines) for individual animal in both NP groups are shown. (C) Total titers against 426.Mod.Core (red circles) and HxB2.WT.Core (blue circles) for all animals are shown; ‘*’ indicates significant differences using Mann-Whitney test. (D) CD4-BS specific values against the indicated proteins are shown; ‘*’ indicates significant differences using two-sample t-test assuming unequal variances. A pool of pre-bleed samples was used as an internal control in all ELISAs.

Irrespective of the NP form of the immunogen, the majority (90-98%) of the autologous anti-426c.Mod.Core antibodies targeted the CD4-BS on that protein, as demonstrated by the lower plasma antibody titers to the CD4-BS knock-out (KO) version of 426c.Mod.Core (Fig 1A: dotted lines and Fig 1D: red squares). Most of the heterologous anti-HxB2.WT.Core antibodies also recognized the CD4-BS on that protein (Fig 1B and 1D), however, a higher proportion of these heterologous CD4-BS antibodies were elicited by the Fer NP form than the C4b NPs (Fig 1D: blue squares). Our results suggest that the valency of the immunogens affect the robustness of the elicited plasma antibody responses.

### Durable autologous and heterologous plasma antibody responses elicited irrespective of the valency of the immunogen

We next examined whether the longevity of the elicited antibody responses was affected by the NP valency. To this end, new groups of animals were immunized with either NP form of immunogen (four animals per group), and the plasma titers of the elicited antibodies were determined over a period of 23 weeks. The autologous plasma antibody titers peaked at week 2 and week 4 for the Fer and C4b groups respectively (Fig 2A: red line). These titers were sustained at high levels over the course of observation in the Fer NP group, but gradually decreased in the C4b NP group; however, there were no statistical differences between the two groups at any time point. In contrast, the relative proportion of the autologous anti-CD4-BS antibody responses slowly decreased during the period of observation in both NP groups (Fig 2B), with week 23 responses being significantly lower than the corresponding peak responses with p<0.05 (Fig 2B: red circles).

**Fig 2.**
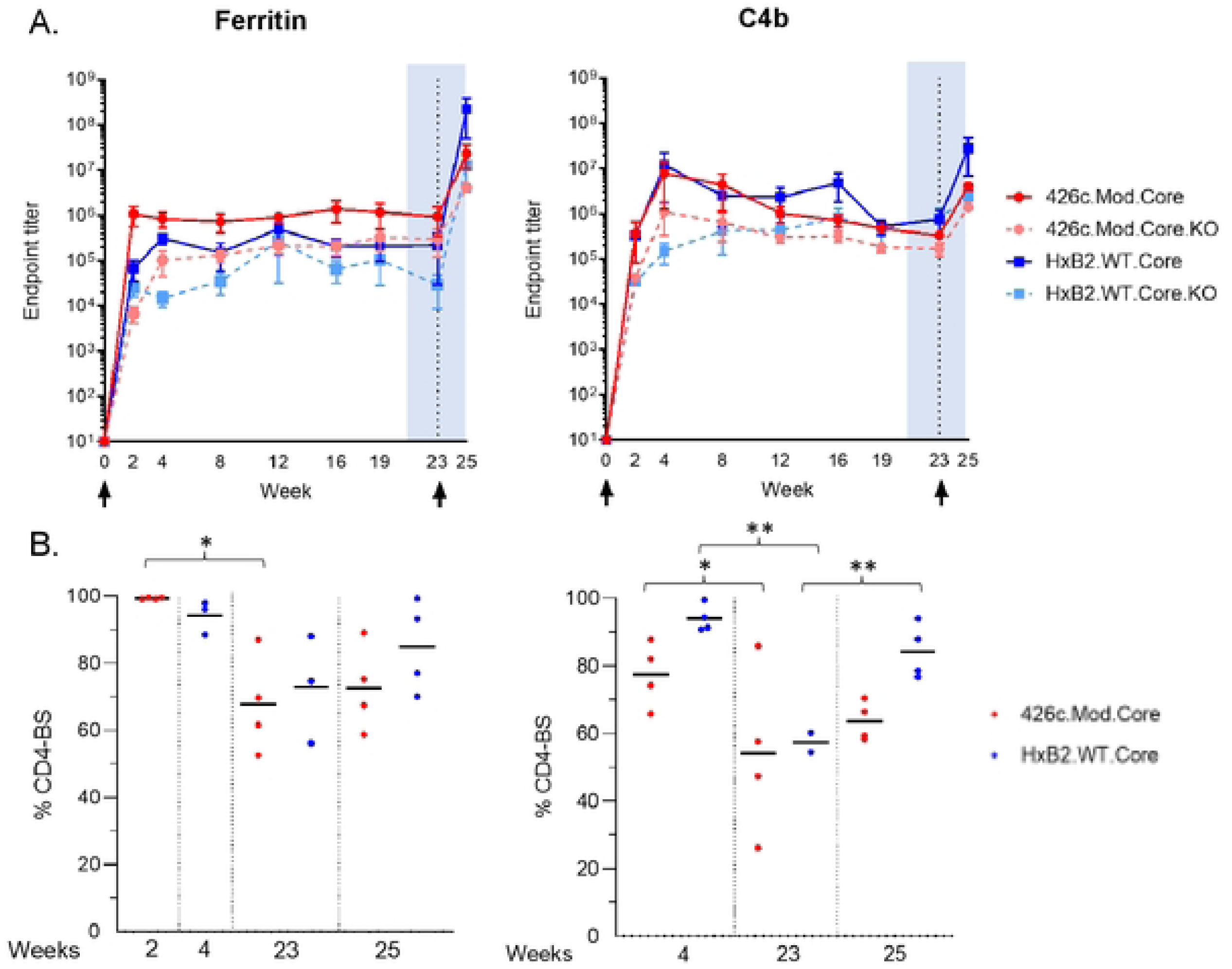
Antibody responses during the course of immunization study. Mice (n = 4) were primed with adjuvanted 426c.Mod.Core-Fer or C4b NPs at week 0, followed by immunization with corresponding NP form of adjuvanted HxB2.WT.Core at week 23. Mice were bled at the indicated time points (x-axis) and plasma was assayed by ELISA for binding. (A) Mean endpoint titers with s.e.m. values against 426.Mod.Core (red solid line), HxB2.WT.Core (blue solid line), as well as their corresponding antigens with CD4-BS knock-out (KO) (dotted lines) are shown. (B) CD4-BS specific percentages against 426.Mod.Core (red circles) and HxB2.WT.Core (blue circles) are shown for indicated time points with ‘*’ indicating significant differences using two-sample t-test assuming unequal variances. A pool of pre-bleed samples was used as an internal control in all ELISAs.

Similarly, the heterologous anti-HxB2.WT.Core antibody responses were durable following immunization with either NP form of 426c.Mod.Core (Fig 2A: blue line), but the fraction of heterologous anti-CD4-BS antibodies gradually decreased in both NP groups (Fig 2B: blue circles). We conclude that a single immunization with the 426c.Mod.Core immunogen (irrespective of its multimeric form), elicits long-lasting autologous and heterologous anti-CD4-BS antibody responses whose relative titers slowly decrease over time, but always remain high.

At week 23, animals were immunized with the corresponding NP form of the heterologous HxB2.WT.Core Env. In contrast to the 426c.Mod.Core, HxB2.WT.Core is fully glycosylated including at positions N276 and N463 (Parks et al. 2019). Two week later (week 25), the proportion of plasma antibodies targeting the CD4-BS of the prime immunogen 426c.Mod.Core, marginally increased in both NP groups, from 68% at week 23 to 73% at week 25 in the Fer group, and from 54% at weeks 23 to 64% at week 25 in the C4b group (Fig 2B: red circles).

Similarly, the proportion of antibodies targeting the CD4-BS of the booster immunogen HxB2.WT.Core, increased in both NP groups between weeks 23 and 25 (73% vs 85% in the Fer group and ∼57% vs 84% in the C4b group) (Fig 2B: blue circles). We conclude that our heterologous immunization (irrespective of its valency) leads to increase in circulating cross-reactive CD4-BS antibodies.

### Impact of antigen valency on the maturation of VRC01-class BCRs

To determine whether the valency of our prime and boost immunogens affects the rate of somatic hyper mutations (SHMs) that are accumulated in Env+ B cells; memory class-switched individual CD4-BS+ B cells were isolated from immunized animals and their VH/VL genes sequenced. We focused our analysis on the B cells that express VRC01-like BCRs.

Two weeks following the prime immunization with 426c.Mod.Core, Env+ B cells expressing VH1-2*02-derived HCs represented ∼90% of the total HCs in both NP groups (Fig 3A and S1A), and the majority of these HCs (66% in the Fer and 59% in the C4b group) expressed a histidine at position 35 in the CDRH1 domain instead of asparagine (H35N, Fig 3B and S1B). The H35N mutation improves the stability of interaction between CDRH1 and CDRH3 on VRC01-class antibodies (Jardine et al. 2015). The majority of the LCs (∼74% in the Fer group, and 65% in the C4b group), expressed 5-aa long CDRL3s (Fig 3C and S1C). Most of these 5-aa CDRL3s-expressing LCs were derived from the mouse κ8-30*01 VL gene (97% in the Fer and 92% in the C4b group; Fig 3D and S1D), consistent with what we and others have previously reported (Knudsen et al. 2022; Parks et al. 2019; Briney et al. 2016; Jardine et al. 2015; Agrawal et al. 2024). Other mLCs expressing 5-aa CDRL3 were also present in both the NP groups, including 12-46*01, 12-44*01, 4-80*01, 4-72*01, 4-53*01, 15-103*01, and 6-25*01 VL genes (Fig 3E).

**Fig 3.**
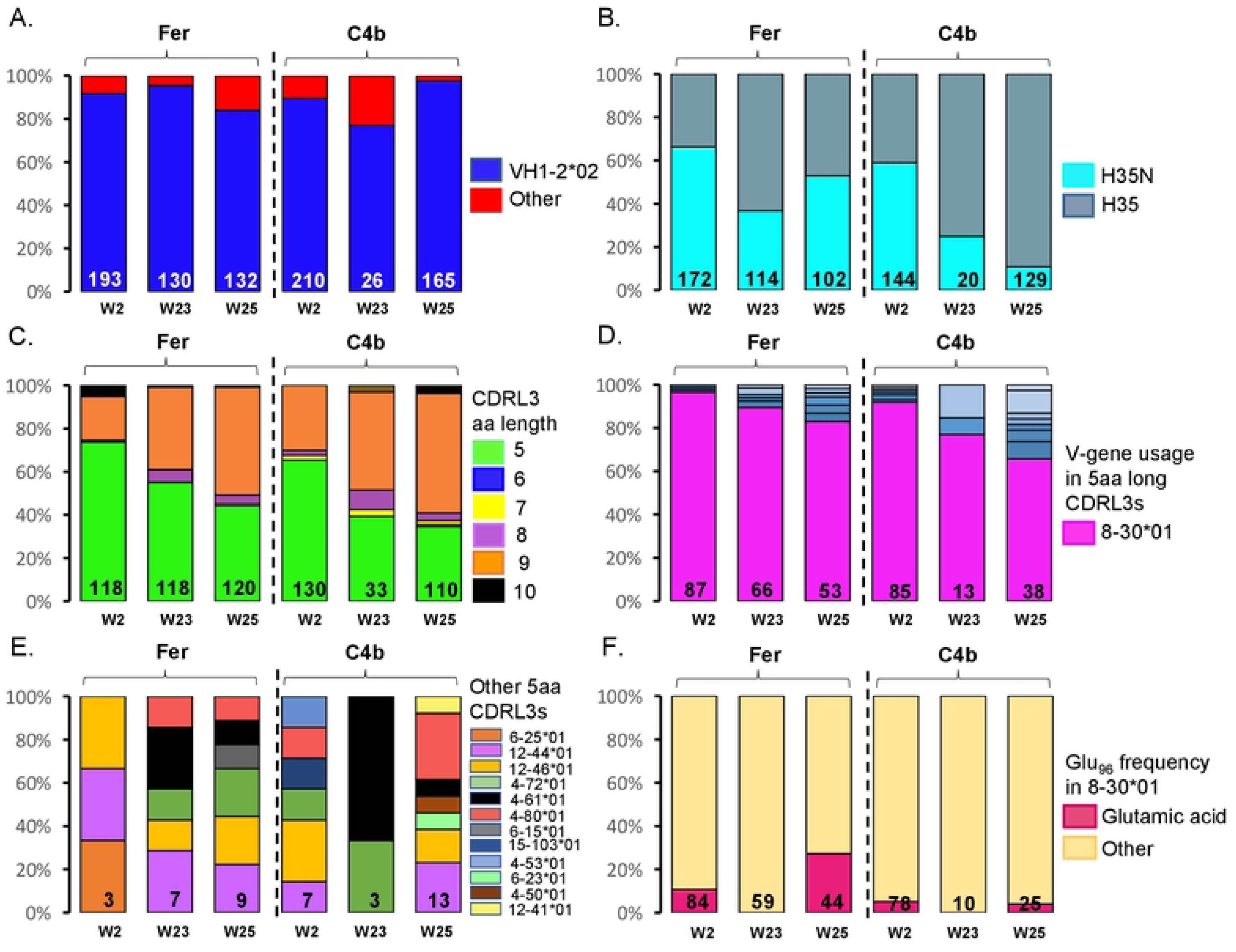
Heavy chain/Light chain sequence analysis after the prime (week 2 and week 23), and boost immunization (week 25). Bar graphs indicate VH (A, B) and VL (B to F) characteristics from individually sorted Env-specific B cells from pooled mouse samples at the indicated time point in each NP group. The number of HC and LC sequences analyzed is shown at the bottom of the bar graph. (A) VH-gene usage, (B) HCs with the H35N mutation, (C) aa length of the CDRL3 domains in the LC, (D) LC-gene usage, where shades of blue slices represent other 5-aa long CDRL3s. (E) LC-gene usage in other 5-aa long CDRL3s and (F), Presence or absence of Glu_96_ within the LC sequences with 5-aa long CDRL3 domains. See also Fig S1.

Furthermore, some of the 5-aa long CDRL3s (∼10% in the Fer and ∼5% in the C4b groups), contained Glu_96_ (Fig 3F, S1E and F); a key feature of mature VRC01-class antibodies (West et al. 2012; Zhou et al. 2013). We conclude that immunization with 426c.Mod.Core, in either NP form, preferentially expands B cells expressing VRC01-like BCRs.

The prevalence of Env+ B cells expressing VH1-2*02 HCs was maintained over time, such that at week 23, >75% of the HC sequences in both groups expressed the VH1-2*02 gene (Fig 3A and S1A). At this time point, ∼37% of VH1-2*02 HCs in the Fer group and ∼25% in the C4b group, expressed the H35N mutation (Fig 3B and S1B). Lower fractions of Env+ B cells expressing LCs with 5-aa long CDRL3 (∼56% in the Fer and ∼40% in the C4b group), were isolated at week 23 compared to week 2 (Fig 3C and S1C). As observed at week 2, majority of those LCs were derived from the mouse κ8-30*01 VL gene (89% in the Fer and 77% in the C4b group; Fig 3D and S1D); but other mLCs expressing 5-aa CDRL3 were also present (Fig 3E and S1D). These results indicate that VRC01-class BCRs that express the desired somatic mutation features are maintained over time following the prime immunization with either NP form of the 426c.Mod.Core germline-targeting immunogen.

B cells expressing VRC01-class BCRs predominated the B cell response following the heterologous HxB2.WT.Core boost immunization in both the NP groups, where ∼84% of HCs in the Fer, and ∼98% of HCs in the C4b groups, were derived from VH1-2*02 (Fig 3A and S1A).

Importantly, the frequency of VH1-2*02 HCs with the H35N mutation significantly increased during the 2-week period after the heterologous immunization (37% at week 23 vs 53% at week 25) in the Fer group only (Fig 3B and S1B).

∼44% of the Env+ B cells in the Fer group and ∼35% in the C4b group, expressed the characteristic 5-aa long CDRL3s of VRC01 antibodies (Fig 3C and S1C). The vast majority of these were still derived from the mouse κ8-30*01 VL gene (83% in the Fer and 66% in the C4b group respectively; Fig 3D and S1D). However, an increase in Env+ B cells expressing LCs with 5-aa CDRL3s derived from other mouse VL genes was evident after the heterologous boost (Fig 3E and S1D). Interestingly, the Glu_96_ LC mutation was also detected in both the groups post boost administration (∼27% in the Fer group and 4% in the C4b group) (Fig 3F and S1E and F). These observations indicate that Env+ B cells with specific VRC01 characteristics are frequent soon after the heterologous boost with either NP form of HxB2.WT.Core Env.

### Differential binding of VRC01-class antibodies isolated following the two NP forms of heterologous Env boost immunization

To directly prove that the VRC01-like BCRs selected by the heterologous boost immunogen, express antibodies with more mature binding and neutralizing properties than the antibodies produced by the BCRs activated by the germline-targeting immunogen alone, we generated mAbs from mice immediately after prime immunization (prime; week 2), right before administration of booster immunogen (prime; week 23), and post boost immunization (boost; week 25), from both NP groups (Table S1). All mAbs expressed the human glVRC01HC paired with mouse κ8-30*01 LC expressing 5-aa long CDRL3; with aa mutations in at least HC or LC (Table S2). The Env-binding properties of these VRC01-class antibodies was then assessed (Fig 4 and 5; Fig S2-4).

**Fig 4.**
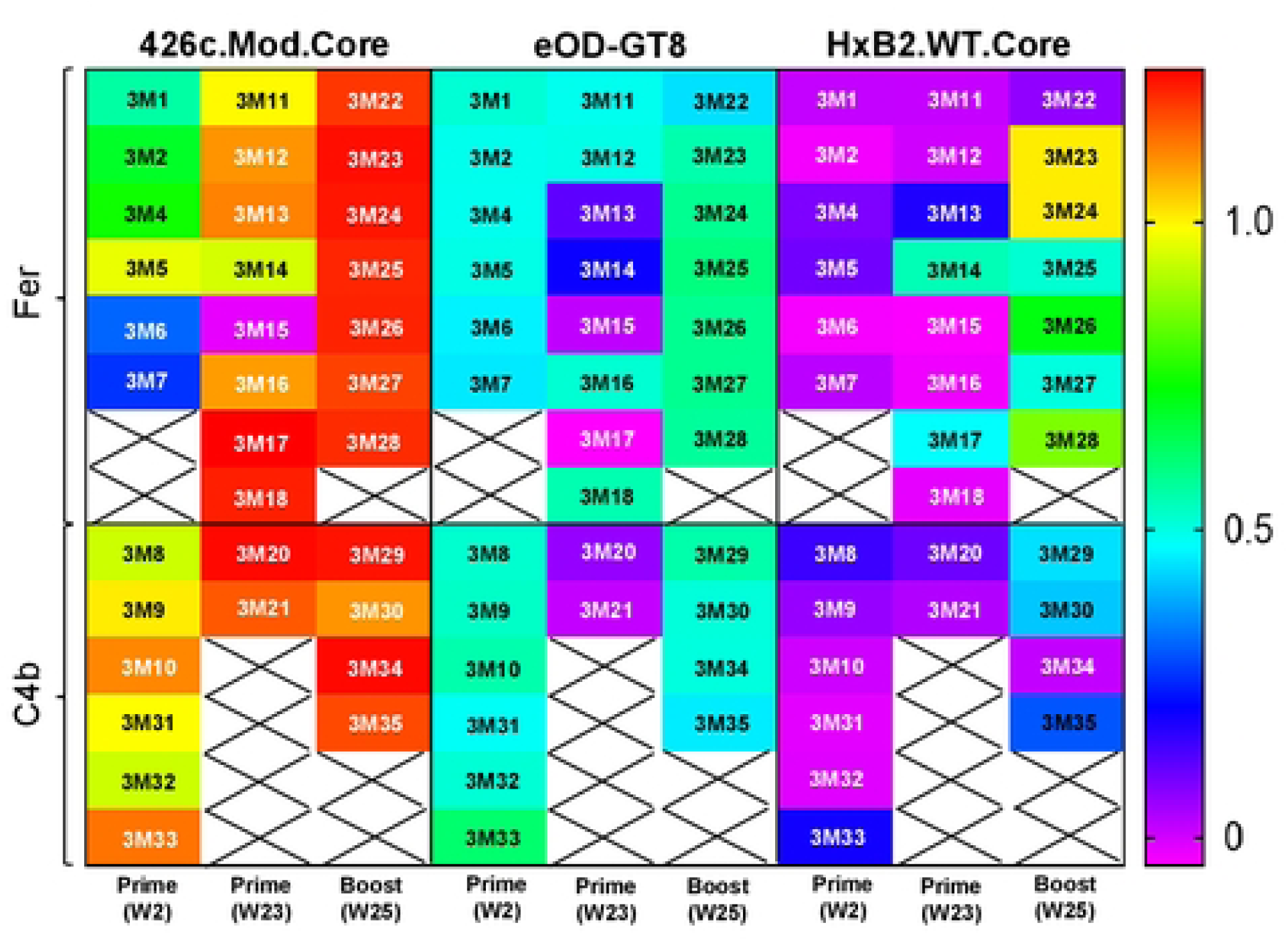
Heat map showing binding properties of VRC01-class mAbs from both NP groups evaluated using BLI assay. 33 VRC01-class mAbs were generated between week 2, week 23, and week 25 timepoints, and tested against the indicated soluble monomeric Envs. Crosses indicate no mAb testing. See also Fig S2 and Table S1.

**Fig 5.**
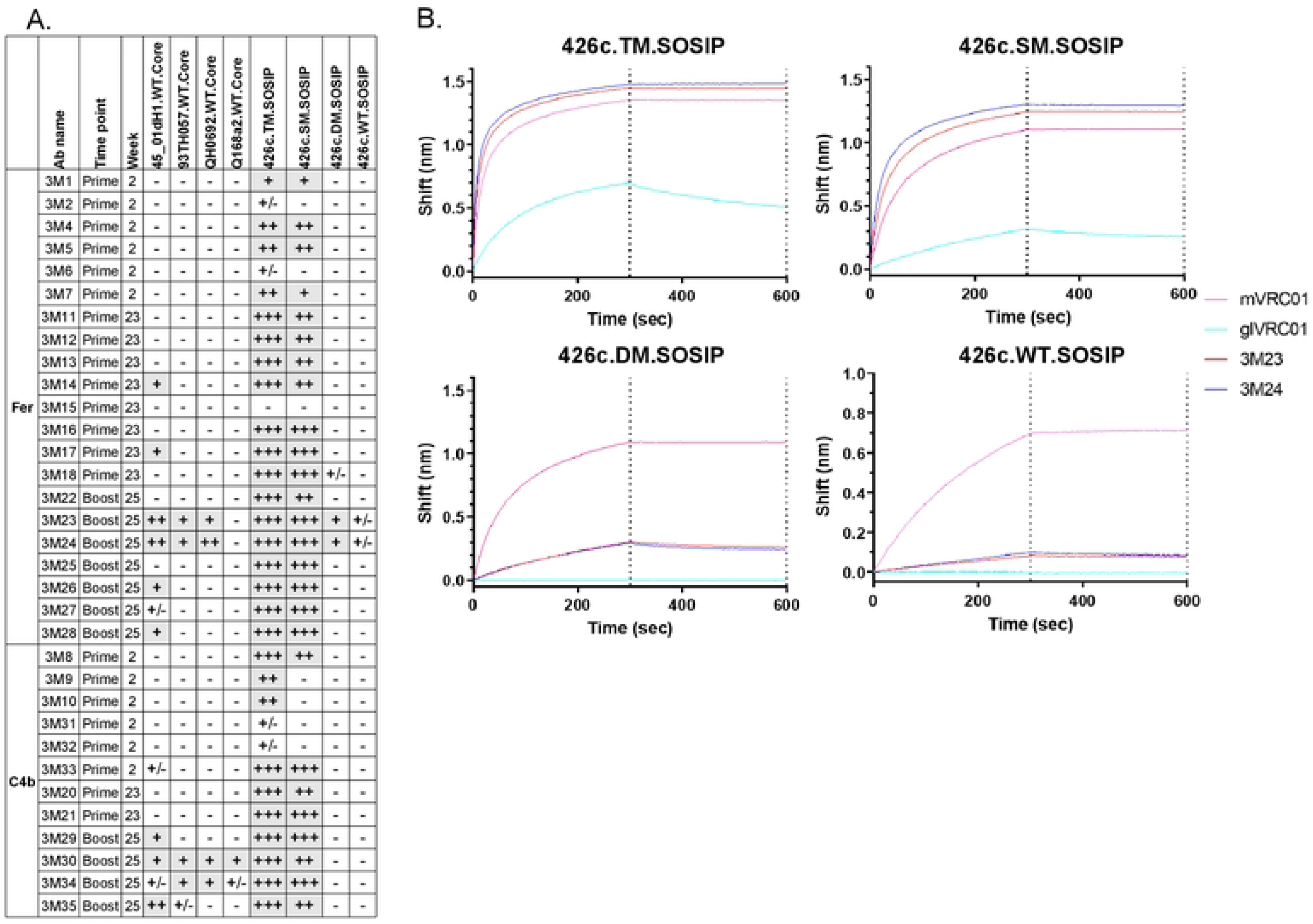
Summary of overall binding properties of VRC01-like mAbs generated at week 2, week 23, and week 25, in both NP groups. (A) 33 VRC01-like mAbs were evaluated against the indicated heterologous WT Core Envs, and variants of 426c SOSIPs. No binding: (-); Up to 0.1: +/-; 0.1 to 0.5: +; 0.5 to 1: ++; and >1: +++. (B) Binding curves of two mAbs (3M23 and 3M24) against the indicated variants of 426c SOSIP are shown. mVRC01 (solid pink line) and glVRC01 (solid cyan line) were included as internal controls in all assays. Black dotted lines indicate end of association and dissociation phases. See also Figs S3 and S4 and Table S1.

Irrespective of the time of sample collection, all mAbs (with the exception of 3M15) recognized 426c.Mod.Core but not its CD4-BS KO version confirming their CD4-BS epitope specificity (Fig 4; binding curves along with that of mVRC01 and glVRC01 mAbs as controls are shown in Fig S2). Similarly, all mAbs, but 3M15, 3M17, and 3M21, bound eOD-GT8 in a VRC01 epitope specific manner (i.e., none bound the CD4-BS KO version of eOD-GT8), with 3M20 showing the weakest binding of all (Fig 4 and S2). Importantly, the post-boost mAbs in both NP groups had faster on rates, slower off rates, and improved overall binding for 426c.Mod.Core. However, only mAbs from the Fer group (post-boost) showed improved binding to eOD-GT8 (Fig S2). Although very few mAbs from weeks 2 or week 23 (i.e., after the prime immunization), bound HxB2.WT.Core, in agreement with our previous observations (Parks et al. 2019), mAbs isolated after the heterologous boost immunization (week 25) displayed improved binding to that protein (and not to its CD4-BS KO version), irrespective of the NP group they were derived from (Fig 4 and S2). These results are in agreement with our previous reports (Knudsen et al. 2022; Parks et al. 2019) that the HxB2.WT.Core selects B cells expressing VRC01-class BCRs that have accumulated mutations enabling them to bypass the N276 and V5 associated glycans on the heterologous HxB2.WT.Core Env.

To determine the level of cross-reactivity of the elicited VRC01-class antibodies, the mAbs were tested for binding to a panel of heterologous WT Cores (Fig 5A; binding curves along with that of mVRC01 mAb, are shown in Fig S3). glVRC01 was also used as an internal control, as it does not recognize Envs with N276-associated glycans (Fig S3, cyan blue line). Several mAbs (3M14, 3M17, 3M23, 3M24, 3M26, 3M28, 3M29, 3M30, and 3M35) bound 45_01dH1.WT.Core; a Clade B Env derived from a virus circulating in patient 45 (Lynch et al. 2015). mAbs 3M23, 3M24, 3M30, and 3M34, also bound to the 93TH057-derived (clade A/E) and QH0692-derived (clade B) WT Core proteins. mAb 3M30 also displayed binding to Q168a2-derived (clade A) WT Core protein (Fig 5A and S3). Importantly, a majority of mAbs that bound the heterologous WT Cores (except 3M14 and 3M17 that were isolated pre-boost at week 23 in the Fer group), were isolated from both NP groups in animals following the boost immunization (Fig 5A and S3); including 3M23, 3M24, and 3M30, that showed the broadest binding.

Next, we examined whether these antibodies could bind the VRC01 epitope on soluble, stabilized Env trimer proteins (SOSIP) (with and without NLGS at position N276 in Loop D and/or in V5). A majority of the mAbs bound both 426c.TM.SOSIP (lacking NLGS at positions N276 in Loop D and N460/N463 in V5) and 426c.SM.SOSIP (lacking only the N276 NLGS) (Fig 5A and B, and Fig S4); indicating that these antibodies can bind in the presence of well-ordered V1-V3 loops not only when the loop D and V5 NLGS are unoccupied (426c.TM.SOSIP) but also when only the loop D N276 glycosylation position is unoccupied (426c.SM.SOSIP). While most of the mAbs were not able to bind to Env trimers that expressed glycans at position N276 (Fig S4), mAbs 3M23 and 3M24 (isolated at week 25 from animals immunized with Fer NP form) showed binding to 426c.DM.SOSIP (N276+/N460-/N463-) (Fig 5A and B). These antibodies also displayed binding (albeit very weak) to the fully glycosylated 426c.WT.SOSIP in this assay (Fig 5A and B). This data along with our previous observation that mAbs 3M23 and 3M24 bind a majority of the heterologous WT Cores, confirms that N276 poses the main obstacle for the maturing VRC01-class antibodies, but also shows that a fraction of the mAbs that are elicited post-booster immunization, are able to partially overcome that obstacle. We conclude that VRC01-class antibodies capable of recognizing the VRC01 epitope on homologous and heterologous Env-derived proteins expressing N276-associated glycans are more effectively elicited by the higher valence nanoparticle form of our immunogens.

### Differential neutralizing potential of the elicited VRC01-class antibodies isolated following the two NP forms of heterologous Env boost immunization

We further examined the neutralizing potential of a subset of these mAbs from both NP groups that were derived from weeks 2, 23, and 25 (n=8). In agreement with the above discussed binding results, none of these mAbs neutralized the 426c.WT virus, irrespective of whether it was produced in 293T or 293 GnTI-cells (GnTI-cells lead to expression of Man5 glycoforms of N-linked glycans that otherwise are processed into large complex-type glycans; Table 1). But all neutralized the TM virus (N276-/N460-/N463-) produced in 293 GnTI-cells in a VRC01-epitope specific manner (as no neutralization was seen against a derivative of TM virus that contains the D279K mutation, which abrogates the neutralizing activity of VRC01-class antibodies). Six of eight mAbs also neutralized the TM virus when expressed in 293T cells (wild type glycans) with post-boost mAbs from both groups neutralizing the virus more potently (Table 1). The mAbs (except 3M35) also neutralized a 426c variant that only lacks the N276 NLGS (SM) when expressed in GnTI-cells and two of eight mAbs (3M23 and 3M24; post-boost mAbs from Fer group) neutralized this virus when expressed in 293T cells as well (Table 1). Importantly, glVRC01 mAb does not neutralize this virus when expressed in 293T cells. Only mAbs 3M23 and 3M24 neutralized two heterologous viruses lacking N276 glycan, when produced in 293 GnTI-cells (Ce703010217_B6.N276Q and CNE55.N276Q) (Table 1); suggesting that additional steric obstacles are present on the heterologous Envs that prevent the binding of these VRC01-class antibodies. Overall, the post-boost mAbs (3M23 and 3M24) isolated from the Fer group, neutralized variants of autologous viruses (when produced in GnTI-cells) more potently and were capable of neutralizing 426c.SM virus (293T cells), and heterologous tier 1b viruses lacking N276 (GnTI-cells) than those isolated from the C4b group. The data strongly suggests that 426c.Mod.Core in either NP form efficiently activates and initiates the maturation of VRC01-class B cells, and administration of corresponding NP form of HxB2.WT.Core as booster immunogen improves the maturation process of these elicited VRC01-class antibodies.

**Table 1.** Neutralizing activities of VRC01-class mAbs against the indicated viruses either grown in 293T or 293S/GnTI-cells. Values represent IC50 concentration in µg/ml. Bold/shaded values indicate samples displaying neutralizing activity. Neutralization IC50 values of these same viruses with the mature VRC01 and germline VRC01 mAb are included for reference. See also Table S1.

|  |  | Prime (W2) |  | Prime (W23) |  | Boost (W25) |  |  |  |  |  |
| --- | --- | --- | --- | --- | --- | --- | --- | --- | --- | --- | --- |
|  |  | Fer | C4b | Fer | C4b | Fer |  |  | C4b |  |  |
|  |  | 3M5 | 3M33 | 3M14 | 3M20 | 3M23 | 3M24 | 3M27 | 3M35 | mVRC01 | glVRC01 |
| 426c | 293T/17 | >50 | >50 | >50 | >50 | >50 | >50 | >50 | >50 | 2.84 | >50 |
|  | 293S/GnTi- | >50 | >50 | >50 | >50 | >50 | >50 | >50 | >50 | 0.12 | >50 |
| 426c.N276D | 293T/17 | >50 | >50 | >50 | >50 | 0.45 | 0.27 | >50 | >50 | 0.41 | >50 |
|  | 293S/GnTi- | 0.78 | 0.054 | 0.061 | 41 | <0.023 | <0.023 | 0.34 | >50 | 0.020 | 0.30 |
| 426c.N276D.N460D.N463D | 293T/17 | >50 | >50 | 3.42 | 2.11 | <0.023 | <0.023 | <0.023 | 0.27 | 0.034 | >50 |
|  | 293S/GnTi- | 0.75 | 0.027 | <0.023 | 0.25 | <0.023 | <0.023 | <0.023 | <0.023 | 0.004 | 0.67 |
| 426c.N276D.N460D.N463D.D279K | 293T/17 | >50 | >50 | >50 | >50 | >50 | >50 | >50 | >50 | >5 | >50 |
|  | 293S/GnTi- | >50 | >50 | >50 | >50 | >50 | >50 | >50 | >50 | >5 | >50 |
| 25710-2.43.N276Q | 293T/17 | >50 | >50 | >50 | >50 | >50 | >50 | >50 | >50 | 0.99 | >50 |
|  | 293S/GnTi- | >50 | >50 | >50 | >50 | >50 | >50 | >50 | >50 | 0.14 | 25.8 |
| Ce1176_A3.N276Q | 293T/17 | >50 | >50 | >50 | >50 | >50 | >50 | >50 | >50 | >5.0 | >50 |
|  | 293S/GnTi- | >50 | >50 | >50 | >50 | >50 | >50 | >50 | >50 | 1.1 | 26.7 |
| Ce703010217_B6.N276Q | 293T/17 | >50 | >50 | >50 | N/A | >50 | >50 | >50 | >50 | 0.086 | >50 |
|  | 293S/GnTi- | >50 | >50 | >50 | N/A | 3.1 | 14.2 | >50 | >50 | 0.003 | 30.1 |
| CNE55.N276Q | 293T/17 | >50 | >50 | >50 | N/A | >50 | >50 | >50 | >50 | 0.100 | >50 |
|  | 293S/GnTi- | >50 | >50 | >50 | >50 | 2.65 | 4.36 | >50 | >50 | 0.018 | >50 |

### Selection of different SHMs by the two NP forms of heterologous Env boost immunogen

Given the observed differences in binding and neutralization potentials of isolated VRC01-class mAbs from the two NP groups, we examined whether these differences were due to increased rates of SHMs in the Fer than C4b groups. The SHM rate with a mean of ∼3.5 and ∼3.8 in the VH1-2*02 HC of the Fer and C4b groups respectively at week 2 was found to be statistically insignificant (Fig 6A). The LCs containing 5-aa long CDRL3 showed a mean SHM rate of ∼3.4 and ∼4.2 in the Fer and C4b groups respectively at week 2 that were also statistically insignificant (Fig 6B).

**Fig 6.**
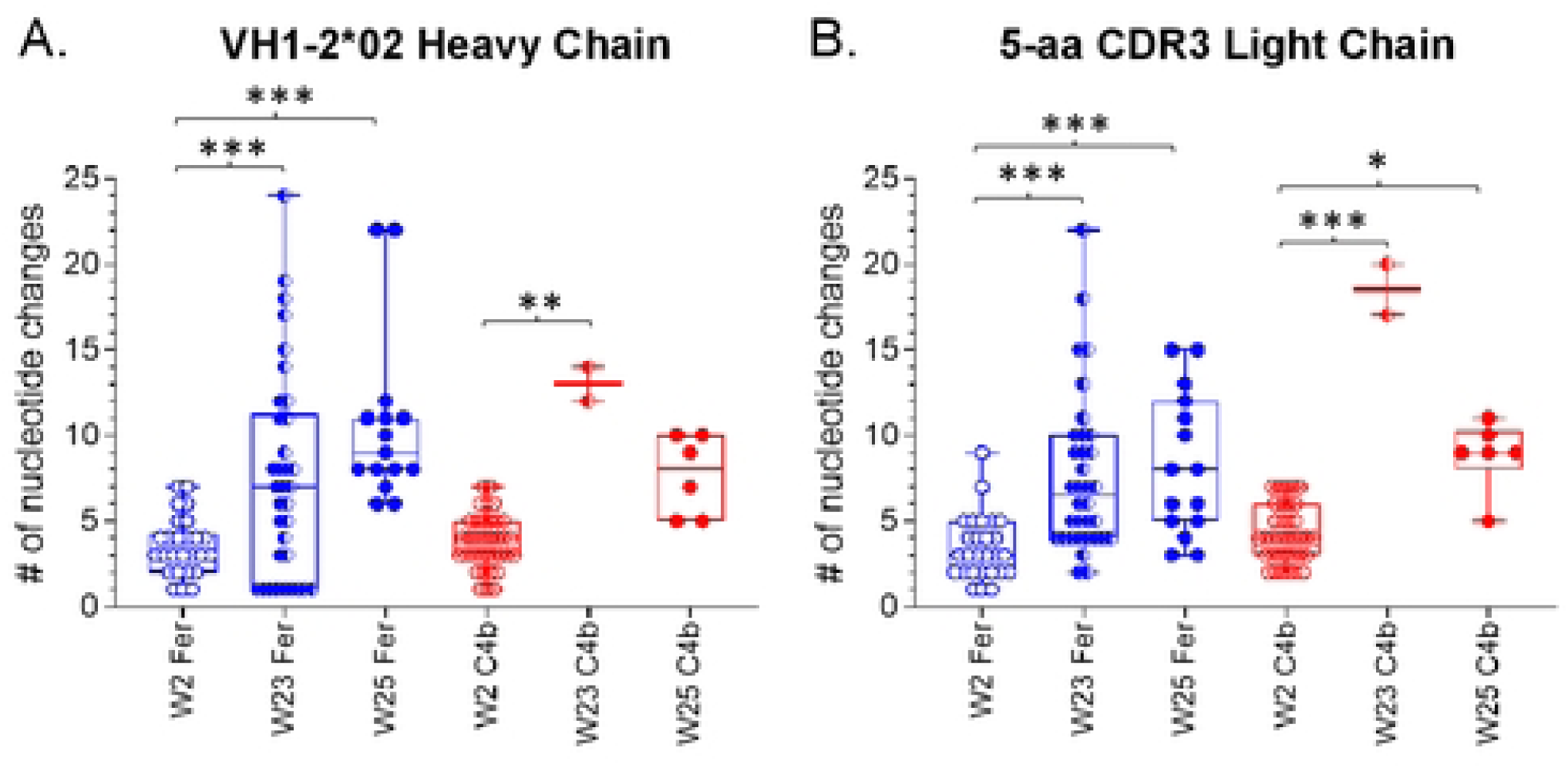
Number of nucleotide changes in the HC and LC of paired sequences at week 2, week 23, and week 25, from both NP groups. Each circle represents a paired sequence and ‘*’ indicates significant differences using Kruskal-Wallis test. See also Fig S5 and Table S1.

Between weeks 2 and 23 the number of nucleotide mutations significantly increased in both the HCs (mean: ∼7.5 for Fer and ∼13 for C4b), and LCs (mean: ∼7.6 for Fer and 18.5 for C4b) (Fig 6A and B) in both groups. However, the SHM rate did not differ significantly between the two NP groups possibly due to a smaller sample size for week 23 in the C4b group. A prolonged GC reaction is evident in animals immunized with either NPs expressing 426c.Mod.Core. The differences in mean SHM rates in both the Fer and C4b groups between weeks 23 and 25 (i.e., 2 weeks post final immunization), were not significant (in either the HC or LC). Thus, the above-described differences in binding and neutralization of the VRC01 antibodies derived following the heterologous boost by the two NP forms, is not due to increased SHMs in the Fer compared to the C4b groups.

We then performed several complementary phylogenetic analyses, to further assess if expansion of particular B cell clones took place between weeks 23 and 25. Intuitively, we suspected that boosting would result in subtrees in which all sequences stemmed from a single timepoint. We thus developed a method for identifying such “single-timepoint” subtrees (see Materials and Methods). In order to assess if we see more of them than we would expect from chance alone, we performed the following randomization procedure: we first constructed a “timepoint-shuffled” sample by shuffling the timepoint labels on our real data in the Fer group. This resulted in a synthetic sample that differs from our real data only in that each point has a randomly chosen incorrect timepoint (a similar approach was used in (Nickle et al. 2003)). We then compared this timepoint-shuffled sample to real data using two metrics (“subtree size” and “subtree distance to ancestor”; Fig S6B), that we designed to highlight any potential subtrees resulting from the booster immunization. “Subtree size” simply measures the size of any such subtree, while “subtree distance to ancestor” is the longest ancestor-to-tip distance in the subtree; where “ancestor” means an ancestral node that has more than one timepoint descending from it. We compared these values in Fig 7 for experimental data (top left) to timepoint-shuffled data (top right). Consistent with the boost induction hypothesis, the large single-timepoint subtrees in experimental data disappeared when we shuffled the timepoints.

**Fig 7.**
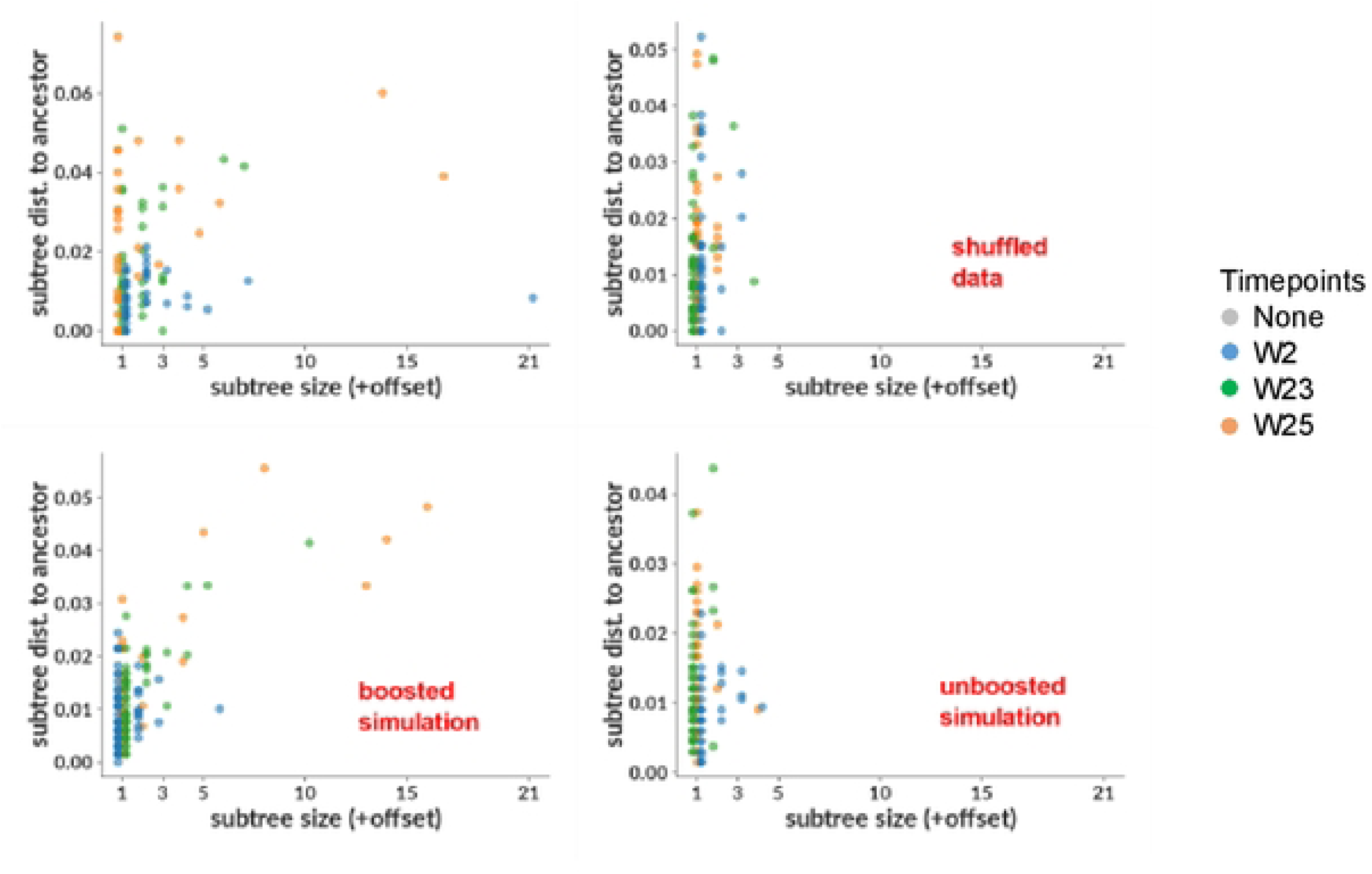
Phylogenetic analysis using HC/LC sequences from the Fer NP group. Size vs distance to common ancestor of single-timepoint subtrees are shown for real data with paired and unpaired sequences (top left), the same real data but with randomly shuffled timepoints (top right); and for boosted simulation (bottom left) and unboosted simulation (bottom right). In both data and simulation, large single-timepoint subtrees occur in cases where we expect to observe the effects of boosting (left column), but they are absent where we do not (right column). The effect in common ancestor distance (y-axis) is less clear than in size (x-axis). See also Fig S6.

Although this disappearance of signal in timepoint-shuffled data shows that that signal depends on the structure of our timepoint labels, we wanted to more directly confirm the effect of the booster immunization. To this end, we used computational methods to construct samples of simulated sequences using new modifications to the simulation method from ((Davidsen and Matsen IV 2018) see Materials and Methods). We mimicked the observed characteristics of real data as closely as possible (such as naive rearrangement properties, mutation rates, and sampled timepoints) for two scenarios: “boosted”, which mimics the expected response to selection by a boosting immunogen, and “unboosted”, mimicking the null hypothesis without any boost effect. We show the subtree metrics described above in Fig 7, for the boosted (bottom left) and unboosted (bottom right) simulation. As expected, we observed large week 25 single-timepoint subtrees in the boosted simulation sample, but not in the unboosted sample. We note that we do not have any compelling theoretical explanation for the much more obvious signal in the subtree size compared to subtree ancestor distance; we include both simply for completeness, since the analysis was performed using both metrics, without any prior expectation as to which would prove most useful. We also show the phylogenetic tree, inferred with IQ-TREE (Minh et al. 2020), from which the preceding analyses were derived in Fig S6A, with each tip colored by timepoint, and expressed mAbs indicated in red. It is evident that the isolated subtrees (the two largest examples are indicated with red boxes in Fig S6A) consist entirely of sequences from the post-boost (orange in color) time points, suggestive of a ‘selection’ effect.

We next examined whether the heterologous HxB2.WT.Core immunogen selected subsets of B cells with specific SHMs that were activated by the prime immunogen 426c.Mod.Core. To this end, we compared the sequences in the VH/VL regions of the elicited mAbs between the two NP groups. Indeed, only post-boost Abs in the Fer NP group showed accumulation of additional amino acid residues (indicated in red ovals in Fig 8 and S7) similar to those present in mature VRC01-class Abs. Overall, we conclude that, in the Fer group of animals, HxB2.WT.Core results in the selection of VRC01-class B cell clones expressing antibodies with more mature VRC01-class sequences.

**Fig 8.**
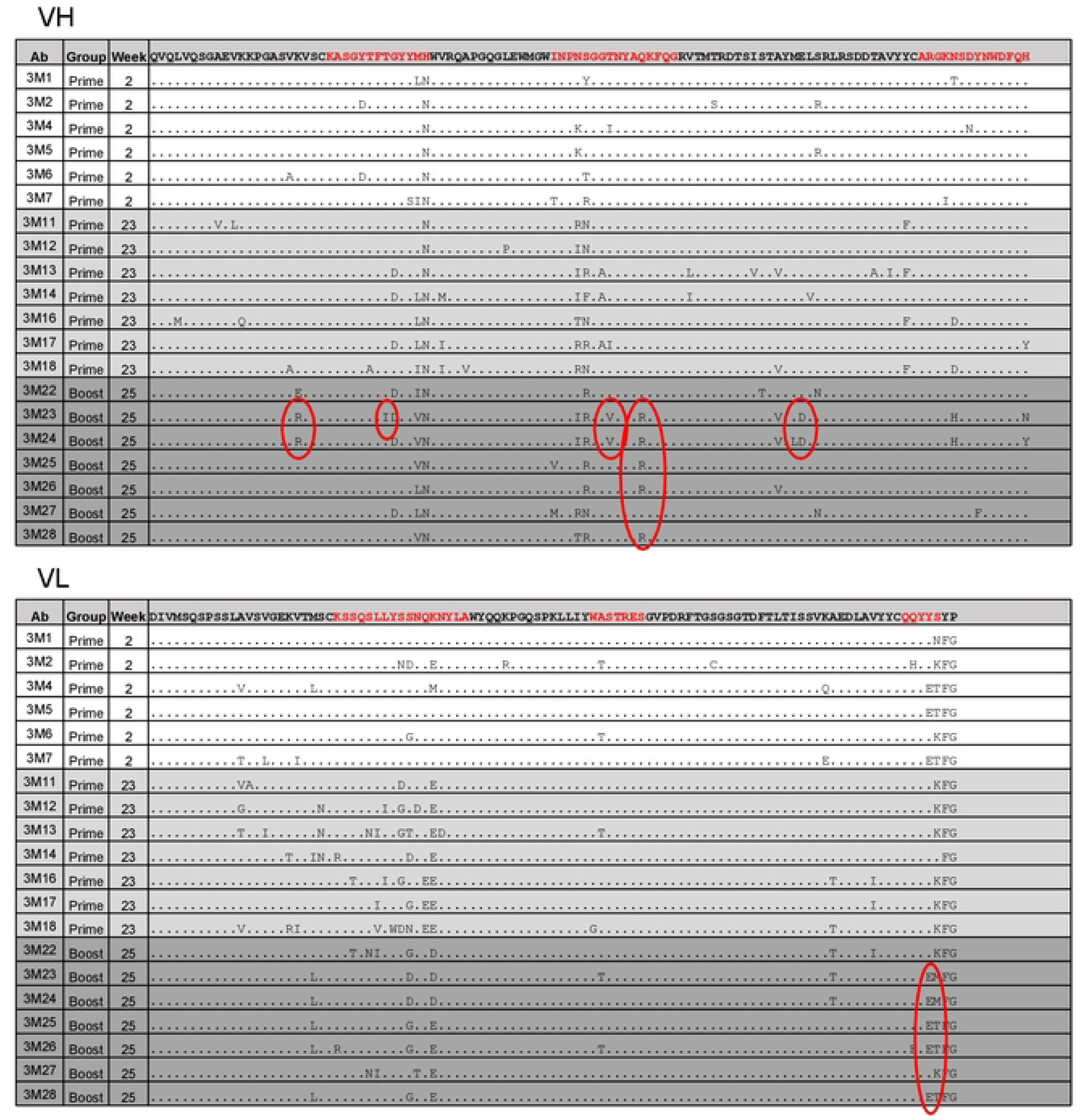
Sequence alignment of VRC01-class mAbs from the Fer NP group. Germline VH1-2*02 and κ8-30*01 sequences are used as reference for alignment, and CDRs are highlighted in red. Red ovals highlight the residues commonly present in mature VRC01-class antibodies. See also Fig S7.

## DISCUSSION

Despite knowledge of the structures of VRC01-class bnAbs, of the importance of specific somatic mutations in defining the broad neutralizing properties of these antibodies, and of the mechanisms of Env-binding and HIV neutralization, the manner by which VRC01-class antibody responses emerge and mature during HIV infection remains poorly understood. In part, this is due to the lack of information on the viral Envs that initiated the activation of naïve B cells expressing glVRC01-class BCRs in those PLWH that developed such responses. In addition, the viral Envs that guided the maturation of VRC01-class antibodies, through the accumulation of somatic mutations at particular positions of their HCs and LCs, remain unknown. In only one PLWH, the concomitant evolution of the viral Env and of VRC01-class HCs and LCs has been evaluated so far, but this evolution of the HCs and LCs was not determined from HC/LC pairs (Umotoy et al. 2019). Thus, natural Envs that initiated and guided the maturation of VRC01-class antibodies in PLWH are not available to be used as immunogens in uninfected persons, in contrast to the case of bnAbs that target epitopes located in the apex region of Env (Andrabi et al. 2015; Landais et al. 2017; Willis et al. 2022) and the VH1-46 lineage of CD4bs bnAbs (Bonsignori et al. 2016). The VRC01-class antibody activation process can be initiated by a single immunization with specifically designed Env-derived germline-targeting immunogens (McGuire et al. 2016; Jardine et al. 2016; Medina-Ramirez et al. 2017), but completion of the maturation process, however, will require multiple booster immunizations with heterologous Envs (Dosenovic et al. 2015; Parks et al. 2019; Tian et al. 2016; Briney et al. 2016; Chen et al. 2021; Agrawal et al. 2024). It is therefore important to identify ways to optimize and accelerate this process.

Our results indicate that the valency of our prime and boost immunogens affect the maturation of the elicited VRC01-class antibody responses. Fer NP immunizations more efficiently lead to the accumulation of somatic mutations that can be found in human VRC01-class antibodies. As a result, the antibodies elicited by the Fer NP immunizations displayed broader Env-binding properties than the antibodies elicited by the C4b NP immunizations. Moreover, the neutralizing activities of mAbs isolated following the prime-boost immunization were not only more potent, but also broader for antibodies elicited in the Fer group of animals (Table 1). Not only did they neutralize 426c-derived viruses more potently, but they also neutralized heterologous viruses lacking N276-associated glycans (Ce703010217_B6 and CNE55), suggesting an early stage of accommodating sequence variability in the core epitope, while also emphasizing the N276 glycan as a major obstacle to overcome. The fact that the potency and breadth of neutralization was higher when the target virus was expressed in 293 GnTI-cells than regular 293T cells, suggests that these mAbs do not yet bind with high enough affinity to virion-associated Envs expressing complex glycans at NLGS surrounding the CD4-BS.

In sum, our study provides direct evidence that the valency of the germline-targeting and 1^st^ heterologous boost immunogen influence the maturation of B cells expressing VRC01-class BCRs. As such, these results are relevant to current and upcoming phase 1 clinical trials that evaluate the ability of germline-targeting immunogens to elicit cross-reactive VRC01-class antibody responses, including those employing 426c.Mod.Core (ClinicalTrials.gov NCT05471076; ClinicalTrials.gov NCT06006546) and upcoming trials combining 426c.Mod.Core and HxB2.WT.Core Envs.

## MATERIALS AND METHODS

### Recombinant HIV-1 envelope protein and tetramer production

Recombinant HIV-1 Env proteins were expressed and purified as previously described (McGuire et al. 2016). The CD4-BS KO version of 426c.Mod.Core contains the D279A, D368R, and E370A mutations whereas the KO version of eOD-GT8 contains the D368R mutation, and substitution of positions 276–279 (DWRD) to NFTA. Self-assembling NPs expressing 426c.Mod.Core and HxB2.WT.Core were produced and purified as previously described (McGuire et al. 2016). They were stored at 4°C for SOSIP NPs and at -20°C for C4b NPs. SOSIP proteins and tetramers of Avi-tagged eOD-GT8, and eOD-GT8.KO, were generated as previously reported (Parks et al. 2019; Lin et al. 2020; Knudsen et al. 2022).

### Mice, immunizations, and sample collection/processing

Transgenic mice expressing the inferred germline HC of the human VRC01 Ab (VRC01^glHC^) and endogenous mouse LCs (Jardine et al. 2015) were bred and kept in house (Animal facility, Fred Hutchinson Cancer Center). Mice were 6–12-week-old at the start of experiments. Env antigens (50μg/mice) and 3M-052-AF + Alum adjuvant were diluted in Tris-NaCl (TBS) and administered intramuscularly with 50μL in each hind leg in the gastrocnemius muscle (total volume 100μL/mouse). Plasma was isolated from blood (collected at indicated time points in tubes containing citrate-phosphate-dextrose solution (CPD; Sigma-Aldrich)), heat inactivated at 56°C and stored short term at 4°C for further analysis. Organs were harvested in cold IMDM media (Gibco), and organ processing for spleens and lymph nodes (LN) was carried out as previously described (Knudsen et al. 2022).

### Enzyme-Linked Immunosorbent Assay (ELISA)

0.1 μM his/avi-tagged proteins (426c.Mod.Core, 426c.Mod.Core.KO, HxB2.WT.Core, and HxB2.WT.Core.KO) diluted in 0.1 M sodium bicarbonate were coated in 384-well ELISA plates (Thermo Fisher Scientific) at room temperature (RT) overnight. After four washes with wash buffer (PBS plus 0.02% Tween20), plates were incubated with block buffer (10% milk, 0.03% Tween20 in PBS) for 1-2 h at 37°C. Post wash step, mouse plasma was added, and serially diluted (1:3) in block buffer. After 1 h of incubation at 37°C, wash step, horse radish peroxidase-conjugated goat anti-mouse IgG (BioLegend) was added and incubated for 1 h at 37°C. After final wash step, SureBlue Reserve TMB Microwell Peroxidase Substrate (KPL Inc.) was added for 5 min. The reaction was stopped with 1 NH_2_SO_4_, and the optical density (OD) was read at 450 nm with a SpectraMax M2 Microplate reader (Molecular Devices). The average OD of blank wells from the same plate were subtracted from all wells before analysis using Prism software.

### Single B-cell sorting and HC/LC V-gene sequencing

Splenocytes or LN cells were stained as previously described (Knudsen et al. 2022), where 1 μM of eOD-GT8, and eOD-GT8.KO, tetramers were used as baits for single cell sorting. Amplification and sequencing of the antibody HC/LC V-genes was performed as previously described (Parks et al. 2019; Lin et al. 2020; Knudsen et al. 2022). Sequences were analyzed using the Geneious software (Biomatters, Ltd.) and the online IMGT/V-QUEST tool (Parks et al. 2019; Lin et al. 2020; Knudsen et al. 2022). SHMs were calculated for sequence length starting from CDR1 to CDR3.

### HC/LC cloning and antibody expression

Gene-specific PCR was carried out using the first round of PCR product to amplify the gene of interest and ligation (Takara Bio) was performed to insert the DNA fragment into human IgG1 vectors: ptt3 for κ LC (Snijder et al. 2018) and PMN 4-341 for γ HC (Scharf et al. 2014). PCR reactions were performed as previously described (Knudsen et al. 2022). Transformation, DNA extraction, and purification was carried out as previously described (Knudsen et al. 2022). Equal amounts of HC and LC DNA were transfected into 293E cells and Abs purified from cell supernatants after 5-7 days using Pierce Protein A agarose beads (Thermo Fisher Scientific).

### Biolayer interferometry

BLI assays were performed on the Octet Red instrument (ForteBio) as previously described (Knudsen et al. 2022; Agrawal et al. 2024). Briefly, anti-human IgG Fc capture biosensors (ForteBio/Sartorius) were used to immobilize mAbs (20 μg/μL), and baseline interference reading measured for 60 s in kinetics buffer (PBS, 0.01% bovine serum albumin, 0.02% Tween-20, 0.005% NaN_3_). Sensors were immersed into wells containing Envs (2 μM) for 300 s (association phase) and another 300 s (dissociation phase). mVRC01 and glVRC01 mAbs were used as internal controls. All measurements were corrected by subtracting the signal obtained from simultaneous tracing of the corresponding Env using an irrelevant IgG Ab. Curve fitting was performed using the Data analysis software (ForteBio).

### TZM-bl neutralization assay

Generated mAbs were tested for neutralization against a panel of selected HIV-1 pseudoviruses using TZM-bl target cells, as previously described (Montefiori 2009). Germline and mature VRC01 mAbs were used as reference in every assay.

### Clonal family and phylogenetic tree inference

Sequences were grouped into clonally related families incorporating heavy/light chain pairing information as described in (Ralph and Matsen IV 2022) with forced over merging (N final clusters set to 1), but otherwise default parameters. The phylogenetic tree was then inferred with IQ-TREE 1.6.12 with default parameters.

### Simulation

We simulated BCR sequences with the bcr-phylo method introduced in (Davidsen and Matsen IV 2018) (updated in (Ralph and Matsen IV 2020)). For this data set, we also added the capability to simulate multiple rounds of GC reactions. To accomplish this, some number of the cells sampled at the end of each GC reaction are selected to start a new GC reaction. The “boosted” simulation samples are then generated with two such GC rounds (where the second round is initiated by the boost vaccination) with week 25 sequences sampled after the second GC reaction. The “unboosted” samples, on the other hand, have only one GC reaction, and week 23 and week 25 sequences are sampled from almost the same pool of sequences (albeit separated by two weeks of evolution). Other simulation parameters were adjusted such that simulation distributions matched those observed in data (results not shown).

### Identification of “boosted” (single-timepoint) subtrees

We expected that the effects of the boost vaccination might be observable in the phylogenetic tree. If working as intended, the boost should stimulate significant new mutation from some subset of existing B cells, which would manifest as new, long branches or subtrees consisting of only sequences from the post-boost time point (week 25). We thus identify such single subtrees by finding all subtrees whose leaves stem from a single timepoint. We quantify these subtrees using two metrics: their size, and the mean distance of their observed nodes from their common ancestor (Fig S6B).

## ACKNOWLEDGEMENTS

This work was supported by grants P01 AI138212, R01 AI143370, and R01 AI177095, from the National Institutes of Health to LS, as well as R01-AI146028 to FAM and HHSN272201800004C to XS. Frederick A. Matsen is an investigator of the Howard Hughes Medical Institute. We thank Access to Advanced Health Institute (AAHI) for providing 3M052-AF+Alum adjuvant used in this study. We thank Translational Research Modeling Services (TRMS) staff members of the Fred Hutchinson Cancer Center for helping with animal work.

---

**Fig S1.**
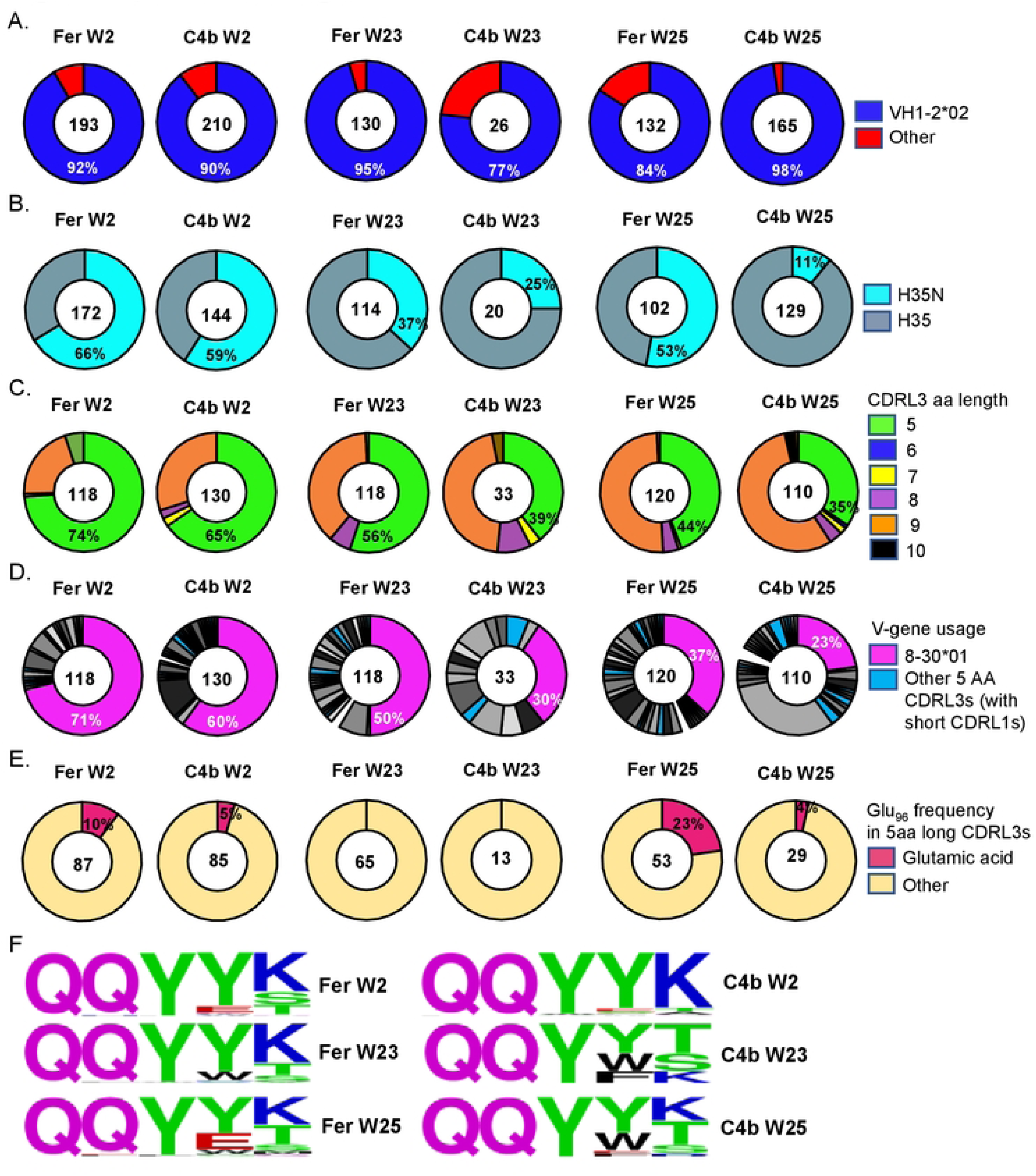
Heavy chain/Light chain sequence analysis at the indicated time points in both NP groups. Pie charts indicate HC (A, B) and LC (C to F) characteristics from individually sorted Env-specific B cells from pooled mouse samples. The number of HC and LC sequences analyzed is shown in the middle of each pie chart. (A) VH-gene usage, (B) HCs with the H35N mutation are shown. (C) aa length of the CDRL3 domains in the LC, (D) LC-gene usage, where shades of grey/black slices represent non 5-aa long CDRL3s and blue indicates other 5-aa CDRL3s. (E) Presence of Glu_96_ within the LC sequences with 5-aa long CDRL3 domains, and (F) Logo plot showing CDRL3 region from the two NP groups at the indicated time points.

**Fig S2.**
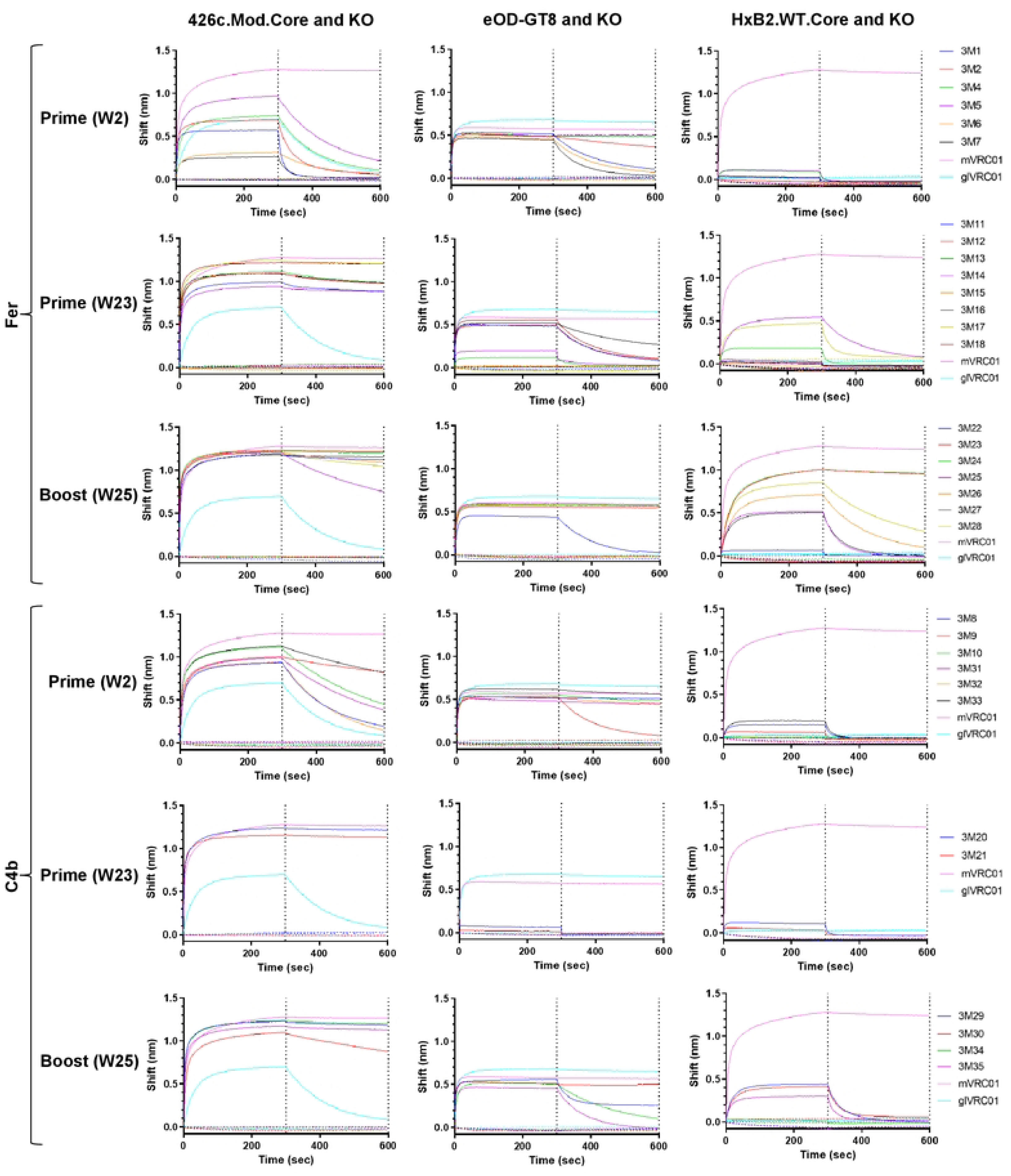
Binding curves of VRC01-class mAbs generated at different timepoints in the two NP groups. mAbs were evaluated against the indicated soluble monomeric Envs and their knock-outs (KO) using BLI assay. mVRC01 (solid pink line) and glVRC01 (solid cyan line) were included as internal controls. Black dotted lines indicate end of association and dissociation phases.

**Fig S3.**
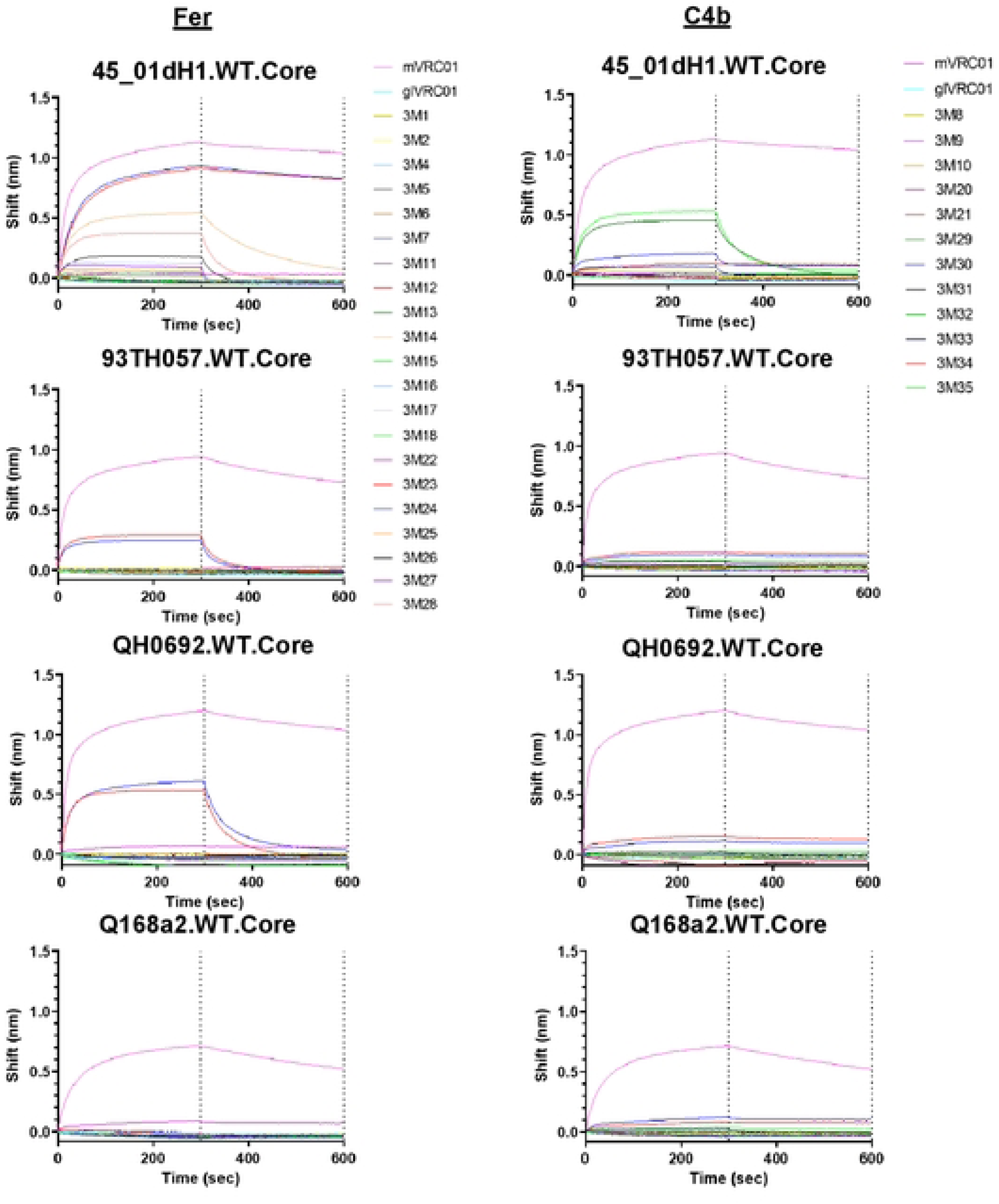
Binding curves of VRC01-class mAbs generated in the two NP groups against heterologous WT.Core Envs. mVRC01 (solid pink line) and glVRC01 (solid cyan line) were included as internal controls. Black dotted lines indicate end of association and dissociation phases.

**Fig S4.**
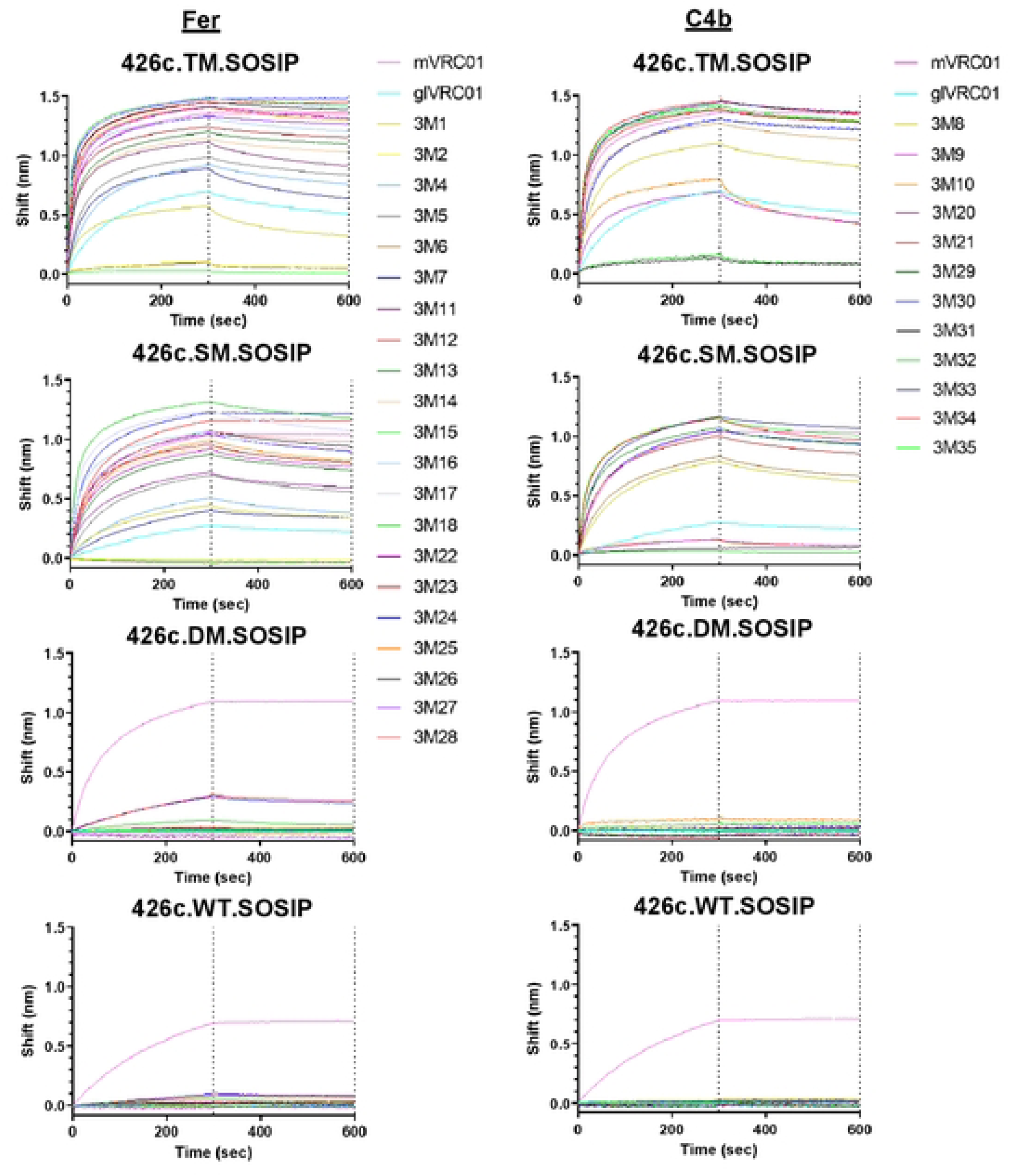
Binding curves of VRC01-class mAbs generated in the two NP groups against indicated variants of 426c SOSIP. mVRC01 (solid pink line) and glVRC01 (solid cyan line) were included as internal controls. Black dotted lines indicate end of association and dissociation phases.

**Fig S5.**
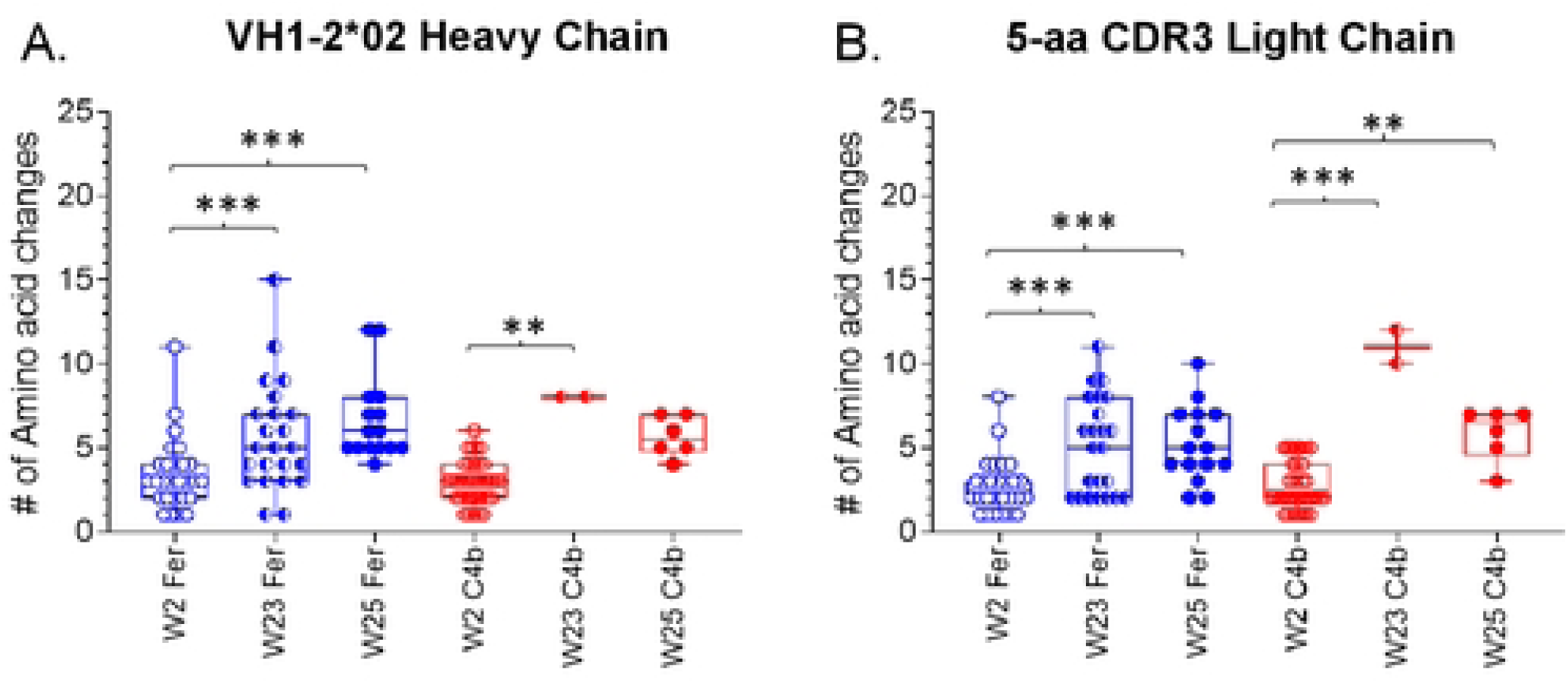
Number of amino acid changes in the HC and LC of paired sequences at week 2, week 23, and week 25, from both NPs groups. Each circle represents a paired sequence and ‘*’ indicates significant differences using Kruskal-Wallis test.

**Fig S6.**
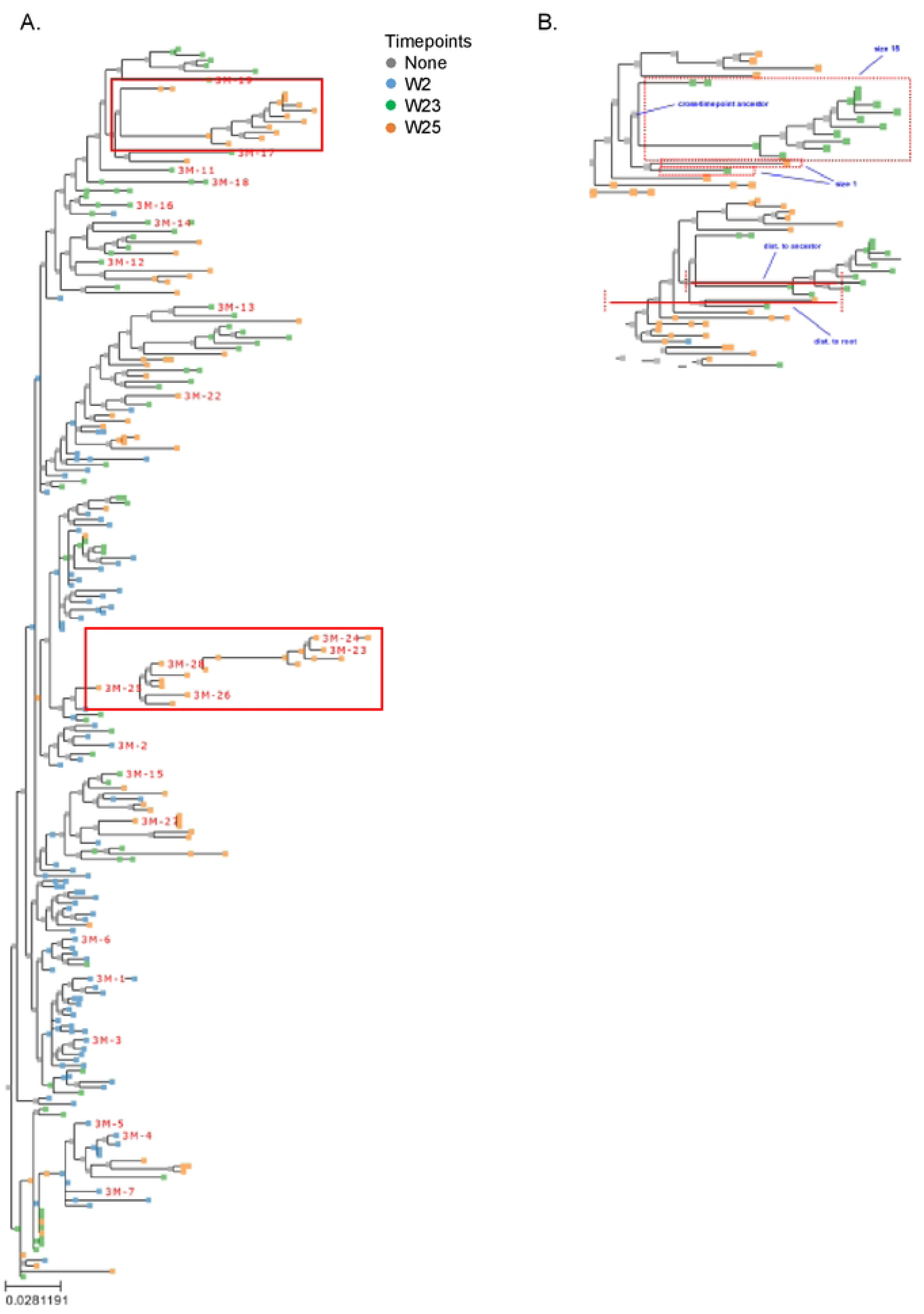
(A) Phylogenetic tree including both paired and unpaired observed sequences in the Fer NP group. Timepoints are colored as indicated (with inferred ancestral sequences in grey), and antibodies chosen for synthesis are labeled in red. (B) Identification of single-timepoint subtrees and calculation of the resulting subtree size (top) and ancestor distance (bottom).

**Fig S7.**
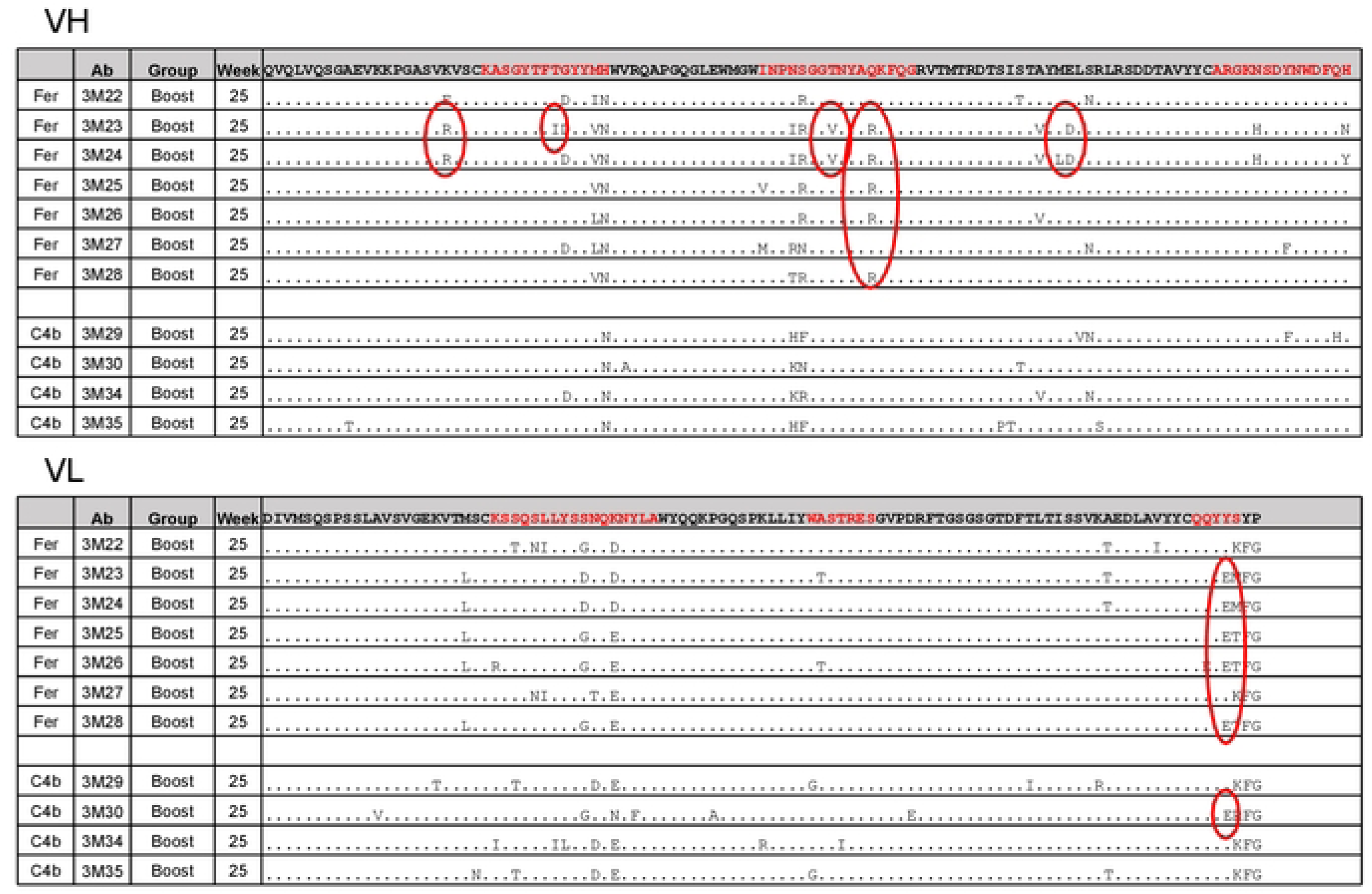
Comparison of sequence alignment of post-boost VRC01-class mAbs from both NP groups. Germline VH1-2*02 and κ8-30*01 sequences are used as reference for alignment, and CDRs are highlighted in red. Red ovals highlight the residues commonly present in mature VRC01-class antibodies.

**Table S1.**
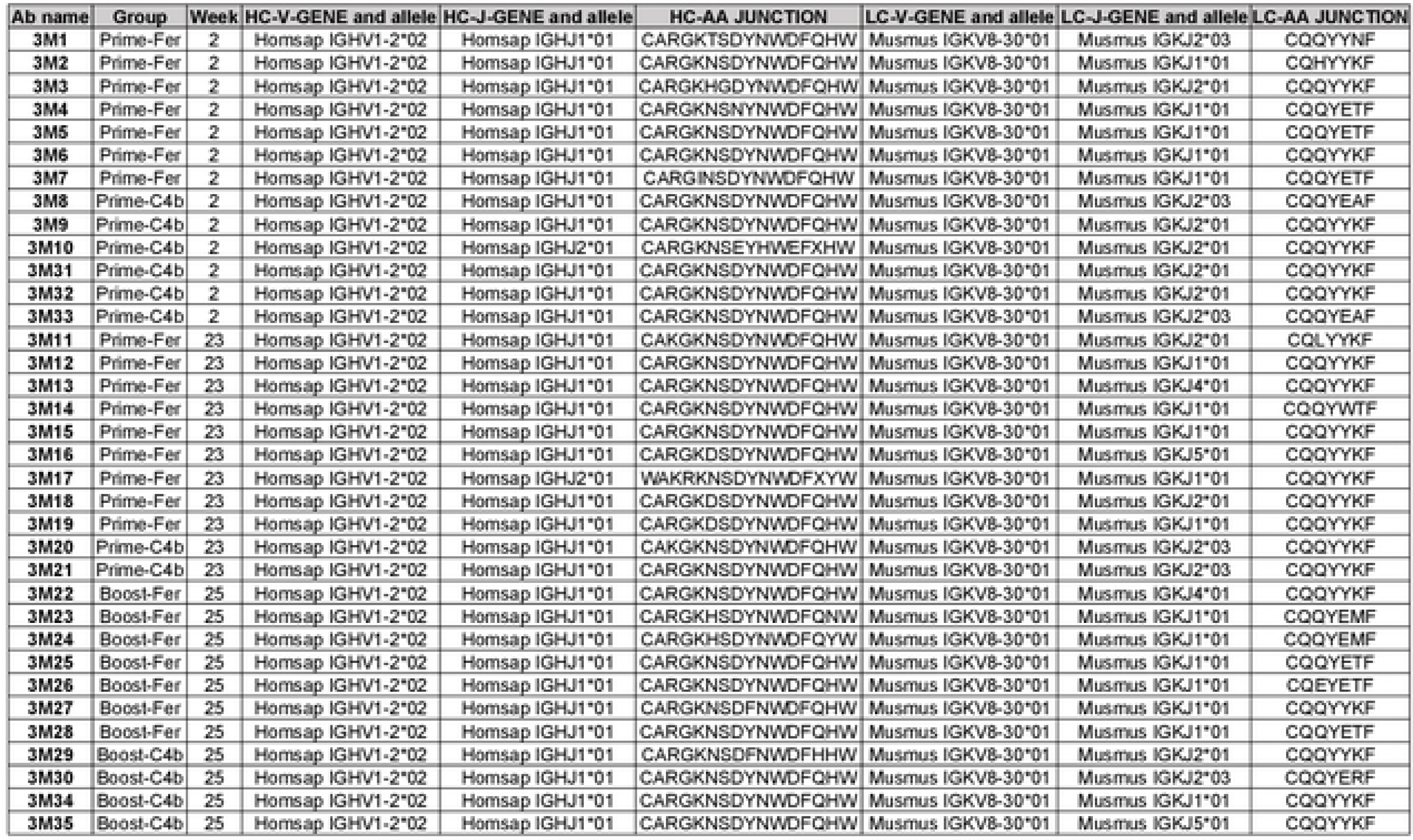
Information on the VRC01-class mAbs isolated at the indicated time points in the two NP groups. A total of 33 VRC01-class mAbs were successfully generated from the immunized animals.

Table S2. HC/LC sequences including that of the VRC01-class antibodies isolated after the final immunization, related to Fig. 4. Amino acid sequences are aligned to the V genes from which they are derived, and CDRs are highlighted in red.

